# Neuronal ketone body utilization couples exercise and time-restricted feeding to cognitive enhancement

**DOI:** 10.64898/2026.02.25.708044

**Authors:** Sebastian F. Salathe, Benjamin A. Kugler, Edziu Franczak, Xin C. Davis, Frederick B. Boakye, Julie Allen, Kyle L. Fulghum, Eric D. Queathem, E. Matthew Morris, Patrycja Puchalska, Peter A. Crawford, John P. Thyfault

**Author notes:** **Co-Corresponding Author:** John P. Thyfault, Ph.D. Professor and Director 3901 Rainbow Boulevard Hemenway Life Sciences Innovation Center, Mailstop 3043 University of Kansas Medical Center Kansas City, KS 66160 Research Scientist Kansas City VA Medical Center Peter A. Crawford, MD, PhD Division of Molecular Medicine, Department of Medicine, 425 East River Rd Minneapolis, MN 55455 University of Minnesota Medical School, Minneapolis, MN, 55455.

## Abstract

Ketogenesis and ketone body metabolism are linked to brain health benefits, including delaying age-related cognitive decline and neurodegeneration. Exercise, particularly when combined with an overnight fast, stimulates ketogenesis and ketone body turnover as well as improves brain metabolism and cognition. Yet, whether ketone metabolism is obligatory for this response is unknown. Here, we use chronic exercise via voluntary wheel running plus time-restricted feeding (VWR+TRF, fasting from ZT10.5-18.5) to explore whether ketone bodies are a potential mediator of exercise-induced brain health benefits in middle-aged mice. To independently distinguish the roles of neuronal ketone body metabolism vs. hepatic ketone body production, we studied middle-age female neuronal-specific SCOT knockout mice and female hepatocyte-specific HMGCS2 knockout mice, respectively. VWR+TRF was compared to sedentary ad-libitum fed (SED+AL) mice to assess the impact on whole-body metabolism (indirect calorimetry), cognition (Barnes Maze and Y-Maze), and molecular adaptations in the hippocampus (proteomics). VWR+TRF robustly upregulated systemic lipid oxidation in all mice, regardless of genotype, during the first 6.5 hours of the dark period. In female SCOT-Neuron-KO mice, we show impaired responses to VWR+TRF in indices of short- and long-term memory. Proteomic analysis of isolated hippocampi revealed that SCOT-Neuron-KO mice failed to globally upregulate key facilitators of synaptic function, including leucine-rich repeated transmembrane proteins, neurexins, and neuroligins. In female HMGCS2-Liver-KO mice, impaired responses to VWR+TRF in indices of short-term memory were paired with an upregulation in ketogenesis machinery in the hippocampal proteome, suggesting potential *in vivo* evidence of cerebral ketogenesis, a mechanism mitigating an otherwise more pronounced behavioral phenotype. Together, these findings suggest that neuronal ketone body utilization is essential, and hepatic ketone production is contributory, to the full cognitive and synaptic adaptations to exercise plus time-restricted feeding, supporting ketone metabolism as a key mechanistic link between metabolic state and brain health in midlife.

## Introduction

Aging is a major risk factor for cognitive decline and neurodegeneration^1–4^. Twelve hallmarks of aging have been identified at the molecular level, including mitochondrial dysfunction, defective macroautophagy, proteostasis impairment, and genomic instability^5^. These deteriorations are particularly relevant in the brain, where highly energetically demanding, predominantly postmitotic neurons rely on mitochondrial energy metabolism and tightly regulated processes to maintain cellular quality control^1,3^. Albeit with inter-individual variability, aging can severely impact the maintenance of quality of life and independent functional status throughout the lifespan^1, 2^. Growing evidence of the plasticity of aging together with the US elderly population projected to nearly double by 2050, emphasizes the need to promote healthy brain aging and delay the onset of cognitive impairment^1, 4, 6^.

Exercise, a modifiable lifestyle factor, has been shown to protect against the functional decline in activities of daily living (ADLs) and to increase healthspan and longevity^6–9^. Increased aerobic fitness has also been demonstrated to protect against age-related reductions in prefrontal, superior parietal, and temporal cortex tissue density in humans^10^. In animal models, regular exercise supports brain health by promoting adaptations that slow age-related cognitive decline, including increased long-term potentiation and neurogenesis in the dentate gyrus, as well as enhanced mitochondrial biogenesis and autophagy^11–14^. While the benefits of exercise continue to be appreciated, the mechanistic mediators of its effects on brain health and function remain largely unknown. Here, we explore ketone bodies as a potential mediator of exercise-induced brain health benefits, whose production can be induced through exercise-mediated liver glycogen depletion and stimulation of fatty acid oxidation, particularly when combined with an overnight fast^15–18^.

The metabolic regulation of ketogenesis and terminal ketone body oxidation has been extensively reviewed by our group^18–20^. Briefly, the primary ketone bodies, acetoacetate (AcAc) and D-β-hydroxybutyrate (D-βOHB), are endogenously produced and classically considered alternative metabolic substrates^18–20^. Conventionally, ketogenesis occurs as adipose tissue lipolysis increases the delivery of free fatty acids to hepatic mitochondria, via carnitine palmitoyltransferase 1 (CPT1), and cyclical rounds of β-oxidation yield acetyl-CoA concentrations that overwhelm citrate synthase activity or oxaloacetate pools^18–20^. Rather than entering the TCA cycle, acetyl-CoAs can instead enter the ketogenic pathway, in which 3-hydroxymethylglutaryl-CoA synthase 2 (HMGCS2) mediates the rate-limiting condensation of acetoacetyl-CoA and acetyl-CoA into HMG-CoA, before being converted to AcAc via HMG-CoA lyase^18–20^. AcAc can either spontaneously decarboxylate into acetone or be further reduced into β-OHB via β-OHB dehydrogenase (BDH1)^18–20^. The liver is the main site of systemic ketone body production because HMGCS2 expression is enriched in hepatocytes^18–20^. While the kidney expresses HMGCS2 and can produce ketone bodies, renal production cannot compensate for the loss of hepatic ketogenesis at a systemic level^21^. Once produced, ketone bodies are exported into the bloodstream and imported into extrahepatic tissues like the brain, via monocarboxylate transporters (MCT), where they undergo terminal oxidation: β-OHB is converted to AcAc through BDH1 and AcAc is converted into acetoacetyl-CoA via succinyl-CoA:3-oxoacid-CoA transferase (SCOT), before being condensed into acetyl-CoA, through mitochondrial thiolase, and entering the TCA cycle^18–20^. Hepatocytes do not express SCOT and are therefore unable to terminally oxidize ketone bodies, while SCOT is expressed in all other metabolic tissues^18–20^. Thus, hepatic lipid metabolism serves as a net producer of ketone bodies that can be metabolized in all other systemic tissues.

Brain energy metabolism accounts for roughly 20% of whole-body energy expenditure^19,22^. Ketones are a significant source of energy in the brain, especially during fasting or carbohydrate restriction (i.e., limited glucose supply)^19, 22^. Neurons express MCT2 and readily use ketone bodies as a fuel source^23^. When SCOT is selectively knocked out in neurons, circulating serum ketone concentrations increase roughly 3-fold and 3.5-fold following a 24-hour and 48-hour fast, respectively, compared to control mice, demonstrating the degree to which neurons consume ketone bodies^24^. Under extreme starvation conditions, ketone bodies can supply up to 60% of the brain’s energetic demand^22^. The brain’s ability to oxidize ketone bodies is currently being leveraged as a therapeutic tool to improve cognition in mild cognitive impairment and Alzheimer’s disease, which are marked by compromised brain glucose metabolism^20, 25–27^. These studies typically employ exogenous ketone body supplementation or ketogenic diets (low-carbohydrate nutritional interventions directed at stimulating ketogenesis)^20, 25–27^. Independent of exercise and beyond their role as metabolic substrates, ketone bodies have potent cellular signaling capabilities^18, 19^. Ketones have been shown to stimulate the production of brain-derived neurotrophic factor (BDNF), activate autophagy, alter post-translational modifications, and protect against oxidative damage in the brain^28–30^.

Exercise, especially when combined with an overnight fast, stimulates robust ketone body turnover (increased ketone production matched with increased ketone utilization), as shown by Féry and Balasse (1983) leveraging stable isotopes in human subjects^17^. Following the cessation of exercise, circulating ketone body concentrations rise, a state termed post-exercise ketosis^16–20, 31^. This response further demonstrates the robust induction of hepatic ketogenesis and high rates of ketone body utilization in metabolic tissues during exercise. Mattson et al. (2018) have proposed that exercise-induced activation of intermittent metabolic switching, oscillating between carbohydrates (glucose) and fatty acid/ketone body fuels, is a primary driving mechanism behind the cognitive benefits of exercise^15^. Taken together, this suggests ketone bodies could mediate exercise-induced brain health benefits. However, *in vivo* approaches mechanistically targeting ketone body metabolism are lacking. Here, we utilized chronic exercise via voluntary wheel running plus time-restricted feeding (VWR+TRF), which stimulates daily hepatic ketogenesis and ketone body turnover, to test whether ketone metabolism mediates exercise-induced brain health benefits in middle-aged mice. To tease out the roles of ketone body utilization and production, we also used female and male neuronal-specific SCOT knockout mice and female hepatocyte-specific HMGCS2 knockout mice. Our findings demonstrate that ketone metabolism indeed influences the cognitive effects of VWR+TRF in mice, and that brain-specific molecular mechanisms are differentially impacted with inhibition of neuronal ketone utilization (SCOT-Neuron-KO) vs hepatic ketone production (HMGCS2-Liver-KO).

## Methods

### Animal Models

Neuronal-specific succinyl-CoA:3-oxoacid-CoA transferase knockout (SCOT-Neuron-KO) mice were generated by crossing *Oxct1^flox/flox^* and *Oxct1^flox/-^* mice with transgenic mice expressing *Synapsin I-Cre* recombinase, all on a C57BL/6 background, as previously described^24^. Littermate *Oxct1^flox/flox^*mice served as controls (Con).

Hepatocyte-specific 3-hydroxymethylglutaryl-CoA synthase 2 knockout (HMGCS2-Liver-KO) mice were generated by crossing *Hmgcs2^flox/flox^* mice with transgenic mice expressing *Albumin-Cre* recombinase, all on a C57BL/6 background, as previously described^32^. Littermate *Hmgcs2^flox/flox^*mice served as controls (Con).

### Experimental Timeline

Female and male (27-31 week old) SCOT-Neuron-KO & Con mice were individually housed and transitioned to an open-source low-fat research diet (LFD; D12110704; 10% kcal fat and 3.5% kcal sucrose). All mice were housed at room temperature (22 °C), on a reverse light cycle (dark: 10:00-22:00), and with *ad libitum* access to water. After 2 weeks of acclimatization to diet and housing conditions, all mice underwent baseline behavior testing, before being grouped by equal bodyweights, but otherwise randomly assigned to two treatments: 1) sedentary plus *ad libitum* fed (SED+AL) or 2) voluntary wheel running plus time-restricted feeding (VWR+TRF, outlined in Figure 1A). While SED+AL mice were housed under standard conditions with unlimited access to food, VWR+TRF mice were given unrestricted access to vertical running wheels (Med Associates) and fasted Monday through Friday (5 days per week) from 8:30 to 16:30 (ZT10.5 to ZT18.5, ZT = zeitgeber time). VWR+TRF mice were given 1 week to adjust to the fasting paradigm before VWR began and running distance was recorded daily, in kilometers per day. Thus, mice had access to wheels and a lack of food 8 hours per day for an 18-week duration. After 12 weeks of intervention, static blood ketone body levels were measured following a tail nick using Precision Xtra blood ketone test strips (Abbott) to demonstrate ketogenic capacity under a 24-hour fast in VWR+TRF mice and a fed state in SED+AL mice. After 2 more weeks, all mice underwent endpoint behavior testing before being maintained in indirect calorimetry chambers (Sable Systems International) for 1 week. Similarly aged (30-34 week old) female HMGCS2-Liver-KO & Con mice completed the same treatment paradigm, except they did not undergo static blood ketone measurements, as ketogenic capacity was previously validated using mass spectrometry^32^. All animal protocols were approved by the Institutional Animal Care and Use Committee at the University of Kansas Medical Center, and all animal use adhered to the National Institutes of Health Guide for the Care and Use of Laboratory Animals.

**Figure 1.**
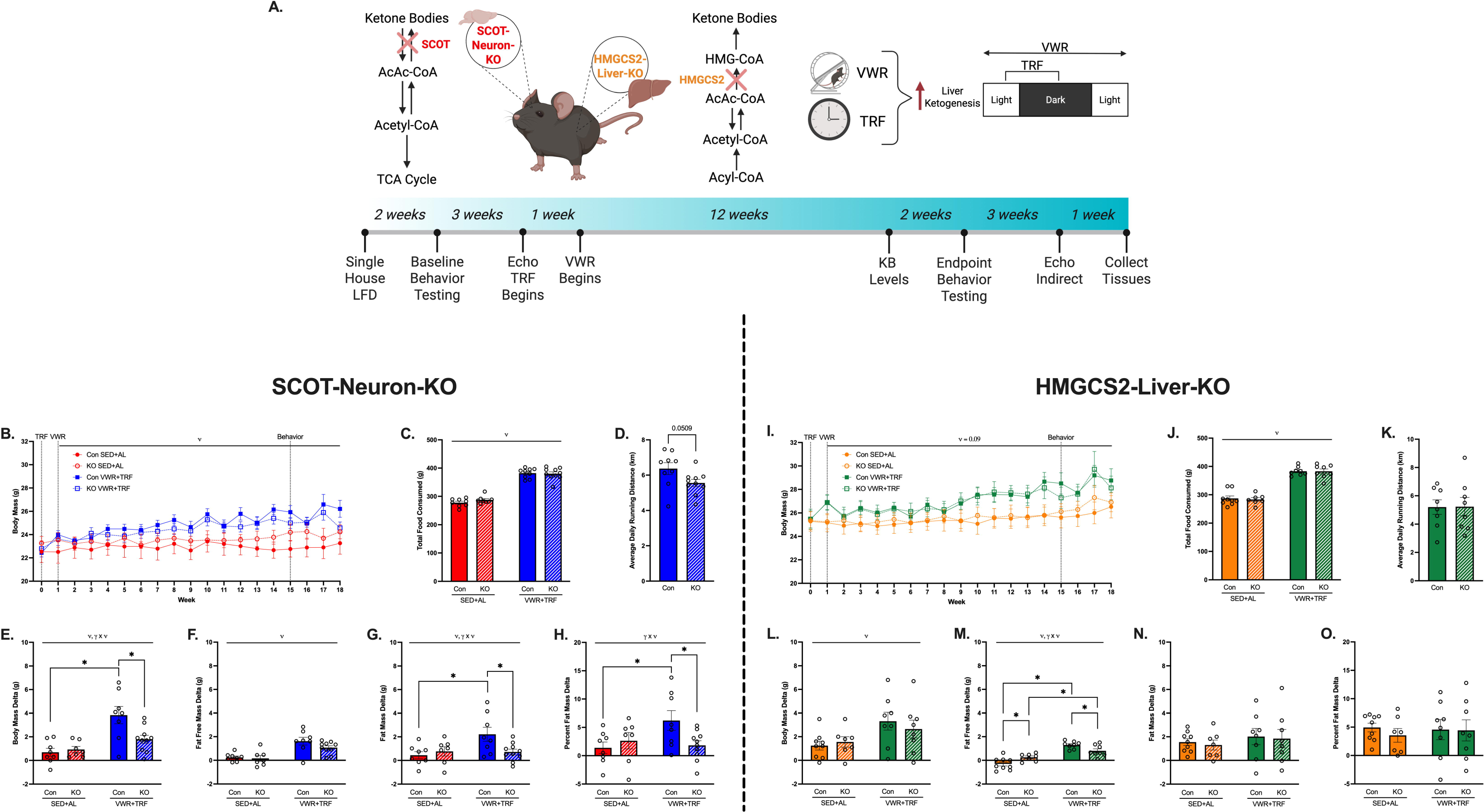
Body mass and composition in SCOT-Neuron-KO and HMGCS2-Liver-Ko female mice. (A) Summary of genotypes, intervention, and study timeline. (B-H) SCOT-Neuron-KO and Con mice. (I-O) HMGCS2-Liver-KO and Con mice. (B/I) Weekly body mass. (C/J) Total food consumed over the duration of the study. (D/K) Average daily running distance. Change from baseline in (E/L) body mass, (F/M) fat free mass, (G/N) fat mass, and (H/O) percent fat mass, calculated by dividing fat mass by body mass. Data are represented as means ± SE. *n* = 7-10 per group. ϖ represents main effect of VWR+TRF following Two-way ANOVA. γ x ϖ indicates significant interaction between genotype and VWR+TRF following Two-way ANOVA, with * denoting significant LSD-post hoc test. BioRender was used to generate figure graphics.

### Longitudinal Behavior Testing

All mice were acclimatized to the behavior testing environment for at least 30 minutes before any behavior testing was conducted. A battery of behavior testing was conducted at baseline and following chronic intervention (endpoint), consisting of Forelimb Grip Strength, Rotarod, modified Y-Maze, and Barnes Maze testing. All testing was completed between 7:00 and 13:00 (ZT9 and ZT15) and food was not restricted for VWR+TRF groups until the completion of each days’ testing.

#### Forelimb Grip Strength

Forelimb grip strength was measured using a digital force gauge (Ametek Chatillon DFE Series), as previously described^33^. Forelimb grip strength was performed by gently raising a mouse toward a horizontal grip bar, allowing it to grip with its forepaws only, and gently pulling horizontally in the opposite direction. The peak tension of 5 consecutive trials was recorded, averaged, and normalized to bodyweight.

#### Rotarod Testing

Rotarod (Med Associates) testing was performed as previously described^34^. All mice were habituated to the Rotarod 24 hours before testing began by being placed on the rotarod for 60 seconds at 4 revolutions per minute (rpm). 3 testing trials with an intertrial interval of 30 minutes were conducted. An accelerating rotarod protocol was utilized, 4 to 40 rpm over 5 minutes. The outcome measure was latency to fall, which was defined by a mouse falling off the rotarod or hanging on to the rotarod and rotating for 2 revolutions, and was an average of the 3 testing trials.

#### Modified Y-Maze

Y-Maze testing was adapted from previously described methods^35^. Each mouse was gently placed at the base of arm A and given 5 minutes to explore the Y-Maze. Arm A was 20 cm long and arms B and C were 15 cm long. The maze was shaped like a true Y, arms B and C were 135° off arm A, and 90° off each other. Arm entries were recorded manually and defined by all 4 limbs crossing the threshold of an arm. The spontaneous alternation index was calculated as previously described, using the following equation^35^: (Alterations ÷ [Total Arm Entries – 2]) x 100.

#### Barnes Maze

Barnes Maze testing was adapted from previously described methods and performed with the Noldus Ethovision XT 17 automated tracking software^36^. The arena consisted of a 92 centimeter diameter white circular table raised 70 centimeters above the floor.

Twenty equally spaced holes were located along the circumference of the table, under one of which was a goal-box (the target hole). Surrounding 3 sides of the table, positioned 6 inches away, were 3 6-foot white plastic walls containing visual cues. Three 1600 lumen lightbulbs were suspended from the plastic walls to provide bright light as the motivating adverse stimulus. The orientation of the arena, visual cues, and location of the goal-box were changed between baseline and endpoint testing. Following every trial, the table was rotated 18 degrees, and the table and goal-box were cleaned with 70% ethanol to minimize intra-maze cues and scents. All mice were habituated to the Barnes Maze 72 hours before testing began, by placing a mouse on the center of the table under a cup for 10 seconds, gently guiding it to the target hole, and allowing it to enter the goal-box. After entering, the target hole was covered for 2 minutes. The spatial acquisition/learning phase (Days 1-4) entailed 4 trials per day for 4 days with an intertrial interval of 15 to 20 minutes. A trial consisted of placing a mouse on the center of the table, under a cup for 10 seconds, and then allowing it 180 seconds to find the target hole and enter the goal-box. If the mouse did not enter within 180 seconds, it was gently guided to the target hole and allowed to enter the goal-box. The target hole was then covered for 1 minute to reinforce behavior. During the intertrial interval, each mouse was placed in its home cage. The primary outcomes were latency to enter the goal-box and distance moved. Memory probes were then performed on Day 5, 24 hours after the last Day 4 trial, and on Day 12. A memory probe was conducted by removing the goal-box from the arena, placing a mouse on the center of the table, under a cup for 10 seconds, and then allowing it 90 seconds to explore. The cumulative time spent in different quadrants was automatically calculated by the Noldus software, in which the target zone was defined as the quadrant containing the target hole in the center (with 2 holes adjacent to the left and 2 holes adjacent to the right).

### Indirect Calorimetry

Whole-body energy metabolism was assessed by housing the mice in a 16-cage Promethion indirect calorimetry system, as previously described (Sable Systems International)^37, 38^. All mice were acclimated to the indirect calorimetry cages for 3 days before collecting data over a 24-hour period. VWR+TRF intervention continued throughout the 4 days in the indirect calorimetry cages, food was restricted from ZT10.5 to ZT18.5 (8:30 to 16:30) and mice had unlimited access to vertical wheels. Average hourly energy expenditure (EE) was calculated using a modified Weir equation (EE (kilocalories per hour) = 60 x [0.003941 x VO2 + 0.001106 x VCO2]). An average hourly respiratory exchange ratio (RER) was calculated using the following equation (RER = VCO2/VO2). Average hourly wheel meters were calculated by multiplying the wheels circumference by the number of wheel revolutions. Measures were analyzed on an hour-by-hour basis or averaged over light and dark cycle periods, allowing for the determination of whole-body metabolism during VWR+TRF exposure or in periods of *ad libitum* feeding and relative inactivity on the wheels.

### Anthropometrics and Tissue collection

Food intake and body mass were measured weekly throughout the study. At baseline and endpoint, body composition was measured via magnetic resonance imaging (EchoMRI). Fat free mass was calculated by subtracting fat mass from total body weight and percent fat mass was calculated by dividing fat mass by body weight and multiplying by 100.

All mice were fasted for 2 hours between 8:00 and 10:00 (ZT10 and ZT12) prior to anesthetization using phenobarbital (0.5mg/g bodyweight) followed by terminal exsanguination. Wheels were pulled 48 hours prior to terminal procedure. Blood was collected via cardiac puncture, allowed to clot at room temperature, and placed on ice for 10 minutes. Serum was then isolated following centrifugation at 7,000 x g (10 min, 4°C). Brains were immediately excised and the hippocampus of the right hemisphere isolated following microdissection on an ice cold mold and flash frozen in liquid nitrogen. Livers were also excised and flash frozen.

### Bulk Hippocampal Proteomics

A subset of female SCOT-Neuron-KO & Con and HMGCS2-Liver-KO & Con right hippocampus samples (n=5 per group) were processed for proteomic analysis. Hippocampi were homogenized in RIPA Lysis Buffer (Thermo Scientific) with protease (EDTA-free protease inhibitor cocktail tablet, Roche) and phosphatase (phosphatase inhibitor cocktail 2 & 3, Sigma) inhibitors. Samples were then submitted to the IDeA National Resource for Quantitative Proteomics. SCOT-Neuron-KO & Con and HMGCS2-Liver-KO & Con groups were processed and analyzed independently as previously described^39–48^:

#### Orbitrap Astral DIA – 40 minutes - CME

Total protein from each sample was reduced, alkylated, and purified by chloroform/methanol extraction prior to digestion with sequencing grade modified porcine trypsin (Promega). Tryptic peptides were then separated by a reverse phase Ion-Opticks-TS analytical column (25 cm x 75 um with 1.7 um C18 resin) supported by an EASY-Spray nano-source and stabilized with a Heater THOR Controller (Ion-Opticks) at 60°C. Peptides were trapped and eluted from a (PepMap Neo, 300um x 5mm Trap) using a Vanquish Neo UHPLC nano system (Thermo Scientific) which kept the samples at 11°C before injection. Peptides were eluted at a flow rate of 0.350uL/min using a 35 min gradient from 98% Buffer A:2% Buffer B to 94.5:5.5 at 0.1 minutes to 56:44 at 27.1 minutes followed by a column wash of 45:55 at 29.7 minutes to 1:99 at 35 minutes followed by equilibration back to 98:2. Eluted peptides were ionized by electrospray (2.5 kV) followed by mass spectrometric analysis on an Orbitrap Astral mass spectrometer (Thermo). Precursor spectra were acquired from 380-980 Th, 240,000 resolution, normalized AGC target 200%, maximum injection time 3 ms. DIA acquisition on the Orbitrap Astral was configured to acquire 199, 3 Th window from 380-980 Th, 25% HCD Collision Energy, normalized AGC target 100%, maximum injection time 3 ms. Fragment (MS2) scan range from 150-2000 Th with an RF Lens (%) set to 40. Buffer A = 0.1% formic acid, 0.5% acetonitrile in water Buffer B = 80% acetonitrile, 20% water, 0.1% formic acid

#### Data Analysis: Spectronaut – VSN

Following data acquisition, data were searched using Spectronaut (Biognosys version 20.4) against the UniProt Mus musculus database (Proteome ID: UP000000589, Taxon ID: 10090, 4^th^ version of 2025) using the directDIA method with an identification precursor and protein q-value cutoff of 1%, generate decoys set to true, the protein inference workflow set to Quant 2.0, inference algorithm set to IDPicker, quantity level set to MS2, cross-run normalization set to false, and the protein grouping quantification set to median peptide and precursor quantity. Fixed Modifications were set to Carbamidomethyl (C) and variable modifications were set to Acetyl (Protein N-term), Oxidation (M). Protein MS2 intensity values were assessed for quality using ProteiNorm^40^. The data was normalized using VSN^45^ and analyzed using proteoDA to perform statistical analysis using Linear Models for Microarray Data (limma) with empirical Bayes (eBayes) smoothing to the standard errors^41, 43^. Ingenuity Pathway Analysis (IPA, Qiagen) was then used to determine up- and down-regulated pathways, within given comparisons, as previously described^49^. Calculated log_2_ Fold Changes (log_2_FC) and p-values were used to determine directionality and Z-scores of each pathway. Pathways with a Z-score greater than or equal to an absolute value of 2 and with a -log_10_p-value greater than 1.3 were considered significant. VSN normalized exclusive intensities were also used to compare protein abundances across all groups as previously described^49^. Within each sample, the VSN normalized exclusive intensity of the protein (or sum of proteins) of interest was normalized to the total summed intensity of that sample (total protein intensity).

### Statistical Analysis

Statistics were performed using Prism 10 (GraphPad). Outliers were identified and removed using the Grubbs method. A two-tailed unpaired t-test was used to assess for differences between KO and Con for baseline behavior testing and average daily running distance. Two-way ANOVA was conducted to determine interactions and main effects of genotype and/or VWR+TRF for baseline body composition, total food consumed, average respiratory exchange ratio (RER), average energy expenditure (EE), changes in body composition, changes in behavior testing outcomes, endpoint spatial learning/acquisition rates, and protein expression. Two-way ANOVA was also conducted to determine interactions and main effects of genotype and/or fasting for average wheel meters. Statistical analysis was conducted within SCOT-Neuron-KO and Cons and within HMGCS2-Liver-KO and Cons, but not across genotypes. Following a significant interaction and/or main effect, a Fisher’s least significant difference post hoc analysis was performed. A repeated measures ANOVA was used in IBM SPPS Statistics 31 to determine changes in body mass over time.

## Results

### SCOT-Neuron-KO and HMGCS2-Liver-KO mice show differentially altered body composition in response to VWR+TRF

A summary of the mouse genotypes, the combined intervention, and the study timeline is shown in Figure 1A. At baseline there were no differences in age (∼35 weeks) or body composition (∼15% body fat) between female SCOT-Neuron-KO & Con mice, female HMGCS2-Liver-KO & Con mice (**Table 1**), or male SCOT-Neuron-KO & Con mice (**Table S1**). VWR+TRF increased the body mass of both female SCOT-Neuron-KO & Con mice (**Fig. 1A**, main effect of VWR+TRF, p<0.05). Male SCOT-Neuron-KOs showed a modest, non-statistically significant increase in bodyweight over Con mice (**Supp. Fig. 1A**, main effect of genotype p=0.097). VWR+TRF increased food intake in both female SCOT-NEURON-KO & Con mice over the course of the study (**Fig. 1C**, main effect of VWR+TRF p<0.0001). Female SCOT-Neuron-KO showed a modest, non-statistically significant decrease in daily running distance compared to Con mice (**Fig. 1D**, p=0.051). Similarly, male SCOT-Neuron-KO & Con mice consumed more food with VWR+TRF (**Supp. Fig. 1B**, main effect of VWR+TRF p<0.0001) and ran the same distance (**Supp. Fig. 1C**). Body mass significantly increased with VWR+TRF in female SCOT-Neuron-KO & Con mice (**Fig. 1E**, main effect of VWR+TRF p=0.0001), but the effect was blunted in SCOT-Neuron-KOs (**Fig. 1E**, genotype and VWR+TRF interaction, p<0.05). This effect on body mass was predominantly driven by differences in fat mass, rather than fat-free mass. As expected, both female SCOT-Neuron-KO & Con mice gain significantly more fat-free mass in response to VWR+TRF (**Fig. 1F**, main effect of VWR+TRF p<0.0001). We identified significant increases in fat mass with VWR+TRF in female SCOT-Neuron-KO & Con mice (**Fig. 1G**, main effect of VWR+TRF p<0.05), but this effect was blunted in SCOT-Neuron-KO mice (**Fig. 1G**, genotype and VWR+TRF interaction, p<0.05). As expected, this interaction also occurred for percent fat mass (**Fig. 1H**, p<0.05). In both male SCOT-Neuron-KO and Con mice, VWR+TRF increased body mass (**Supp. Fig. 1D**, main effect of VWR+TRF p<0.05), an effect predominantly driven by increases in fat-free mass (**Supp. Fig. 1E**, main effect of VWR+TRF p<0.0001). Male SCOT-Neuron-KO mice showed reduced levels of fat mass compared to Con mice (**Supp. Fig. 1F**, main effect of genotype p<0.05), an effect that did not persist when quantifying fat mass as percent change (**Supp. Fig. 1G**).

**Table 1.** Baseline Anthropometric Data.

| Table 1. Baseline Anthropometric Data |  |  |  |  |  |  |  |
| --- | --- | --- | --- | --- | --- | --- | --- |
|  | Control Mice |  | SCOT-Neuron-KO Mice |  | Two-Way ANOVA p-value |  |  |
|  | SED+AL | VWR+TRF | SED+AL | VWR+TRF | Interaction | Genotype | Intervention |
| **Age at Start of Intervention** |  |  |  |  |  |  |  |
| Weeks Old | 34.79 ± 0.31 | 34.30 ± 0.31 | 34.45 ± 0.31 | 34.24 ± 0.30 | 0.65 | 0.53 | 0.27 |
| **Anthropometric Phenotyping** |  |  |  |  |  |  |  |
| Body Mass (g) | 22.58 ± 0.91 | 22.47 ± 0.35 | 23.26 ± 0.58 | 22.79 ± 0.48 | 0.76 | 0.40 | 0.63 |
| Fat Mass (g) | 3.62 ± 0.47 | 3.23 ± 0.25 | 4.15 ± 0.53 | 3.68 ± 0.49 | 0.92 | 0.29 | 0.35 |
| Fat Free Mass (g) | 18.96 ± 0.46 | 19.24 ± 0.22 | 19.12 ± 0.30 | 19.11 ± 0.39 | 0.69 | 0.97 | 0.70 |
|  | Control Mice |  | HMGCS2-Liver-KO Mice |  | Two-Way ANOVA p-value |  |  |
|  | SED+AL | VWR+TRF | SED+AL | VWR+TRF | Interaction | Genotype | Intervention |
| **Age at Start of Intervention** |  |  |  |  |  |  |  |
| Weeks Old | 37.55 ± 0.39 | 37.07 ± 0.44 | 36.88 ± 0.40 | 36.46 ± 0.24 | 0.93 | 0.10 | 0.24 |
| **Anthropometric Phenotyping** |  |  |  |  |  |  |  |
| Body Mass (g) | 25.28 ± 0.83 | 25.45 ± 0.73 | 25.34 ± 0.76 | 25.48 ± 0.86 | 0.98 | 0.96 | 0.85 |
| Fat Mass (g) | 4.77 ± 0.50 | 5.16 ± 0.48 | 5.71 ± 0.54 | 5.40 ± 0.66 | 0.53 | 0.29 | 0.94 |
| Fat Free Mass (g) | 20.51 ± 0.44 | 20.29 ± 0.38 | 19.63 ± 0.29 | 20.07 ± 0.38 | 0.39 | 0.17 | 0.78 |
| Data are presented as Mean ± SE. n = 7-10 per group. Abbreviations: SCOT, succinyl-CoA:3-oxoacid-CoA transferase; HMGCS2, 3-hydroxymethylglutaryl-CoA synthase; SED+AL, sedentary plus ad libitum; VWR+TRF, voluntary wheel running plus time restricted feeding |  |  |  |  |  |  |  |

In female HMGCS2-Liver-KO & Cons, VWR+TRF tended to increase body mass over time (**Fig. 1I**, main effect of VWR+TRF p=0.09). Female HMGCS2-Liver-KO & Con mice both consumed more food with VWR+TRF (**Fig. 1J**, main effect of VWR+TRF p<0.0001) and ran the same distance (**Fig. 1K**). VWR+TRF increased body mass in both female HMGCS2-Liver-KO & Con mice (**Fig. 1L**, main effect of VWR+TRF p<0.05) and this effect was largely driven by VWR+TRF increasing fat-free mass (**Fig. 1M**, main effect of VWR+TRF p<0.0001). However, the impact of VWR+TRF on fat-free mass was significantly blunted in HMGCS2-Liver-KOs (**Fig. 1M**, VWR+TRF and genotype interaction p<0.01). In SED+AL conditions, female Cons displayed reduced fat-free mass compared to female HMGCS2-Liver-KOs (p<0.05), while in VWR+TRF conditions, female Cons exhibited increased fat-free mass compared to female HMGCS2-Liver-KOs (p<0.05). There were no differences in changes in fat mass or percent fat mass between female HMGCS2-Liver-KO & Con mice (**Fig. 1N-O**).

### VWR+TRF shifts systemic metabolism to fat derived substrates during the fasting period

To determine the requirements of neuronal ketone body oxidation and hepatic ketogenesis for the adaptations in energy expenditure to voluntary wheel running plus time-restricted feeding, we performed indirect calorimetry. Data are shown temporally on an hour-by-hour basis and are also quantified within the dark (ZT12 to ZT24) and light period (ZT0 to ZT12). During the dark period, data are further divided into fasting (ZT12 to ZT18.5) and feeding (ZT18.5 to ZT0) periods. Consistent with the rest of the study, we observed no genotype-dependent differences in running distance in female (**Fig. 2A-B**) or male (**Supp. Fig. 2A-B**) SCOT-Neuron-KO mice. Importantly, in support of our combined intervention, female (**Fig. 2A-B**) and male (**Supp. Fig. 2A-B**) SCOT-Neuron-KO & Con mice undergoing VWR+TRF, were active on running wheels during the fasting period of the dark cycle. As expected, the combination of VWR and TRF markedly decreased RER during the fasting period of the dark cycle in female (**Fig. 2C-D**, main effect of VWR+TRF p<0.0001) and male (**Supp. Fig. 2C-D**, main effect of VWR+TRF p<0.0001) SCOT-Neuron-KO & Con mice, compared to SED+AL, signifying a shift to fat-derived sources, including ketone bodies. Moreover, a 24 hour fast provoked greater circulating D-βOHB concentrations in VWR+TRF mice over SED+AL mice in both females (**Supp. Fig. 3A**, main effect of VWR+TRF p<0.0001) and males (**Supp. Fig. 3B**, main effect of VWR+TRF p<0.0001), with no genotype-dependent effects observed. Concurrently, VWR+TRF increased energy expenditure (EE) during the fasting period of the dark cycle in both female (**Fig. 2E-F**, main effect of VWR+TRF p<0.0001) and male (**Supp. Fig. 2E-F**, main effect of VWR+TRF p<0.05) SCOT-Neuron-KO & Con mice compared to SED+AL. Once food was returned, both female (**Fig. 2A-B**, main effect of fasting p<0.001) and male (**Supp. Fig. 2A-B**, main effect of fasting p< 0.01) SCOT-Neuron-KO & Con mice undergoing VWR+TRF ran less during the remainder of the dark cycle than when fasted. This was likely due to the mice spending time eating, which is supported by a marked increase in RER in female (**Fig. 2C-D**, main effect of fasting p<0.0001) and male (**Supp. Fig. 2C-D**, main effect of fasting p<0.0001) SCOT-Neuron-KO & Con mice undergoing VWR+TRF. This was further supported as EE also increased, likely due to the thermogenic effect of food, when food was returned during the dark period within VWR+TRF groups in both female (**Fig. 2E-F**, main effect of fasting p<0.01) and male (**Supp. Fig. 2E-F**, main effect of fasting p<0.01) SCOT-Neuron-KO & Con mice. During the fed period of the dark cycle, VWR+TRF increased RER (**Fig. 2C-D**, main effect of VWR+TRF p<0.05) and EE (**Fig. 2E-F**, main effect of VWR+TRF p<0.0001) in female SCOT-Neuron-KO & Con mice compared to SED+AL. The same effect occurred for RER and EE in male SCOT-Neuron-KO & Con mice (**Supp. Fig. 2C-D**, main effect of VWR+TRF p<0.01 and **Supp. Fig. 2E-F**, main effect of VWR+TRF p<0.0001). However, male SCOT-Neuron-KOs had reduced EE during the fed period of the dark cycle compared to Con mice, regardless of intervention (**Supp. Fig. 2E-F**, main effect of genotype p<0.05). Interestingly, VWR+TRF sustained an increased RER in female (**Fig. 2C-D**, main effect of VWR+TRF p<0.0001) and male (**Supp. Fig. 2C-D**, main effect of VWR+TRF p<0.0001) SCOT-Neuron-KO & Con mice during the light cycle. In a similar fashion, VWR+TRF also sustained an increased EE during the light cycle in both females (**Fig. 2E-F**, main effect of VWR+TRF p=0.0001) and males (**Supp. Fig. 2E-F**, main effect of VWR+TRF p<0.01). Taken together, VWR+TRF in SCOT-Neuron-KO paired ketogenesis with increased energy expenditure, stimulating robust ketone body turnover, with no genotype-dependent effects.

**Figure 2.**
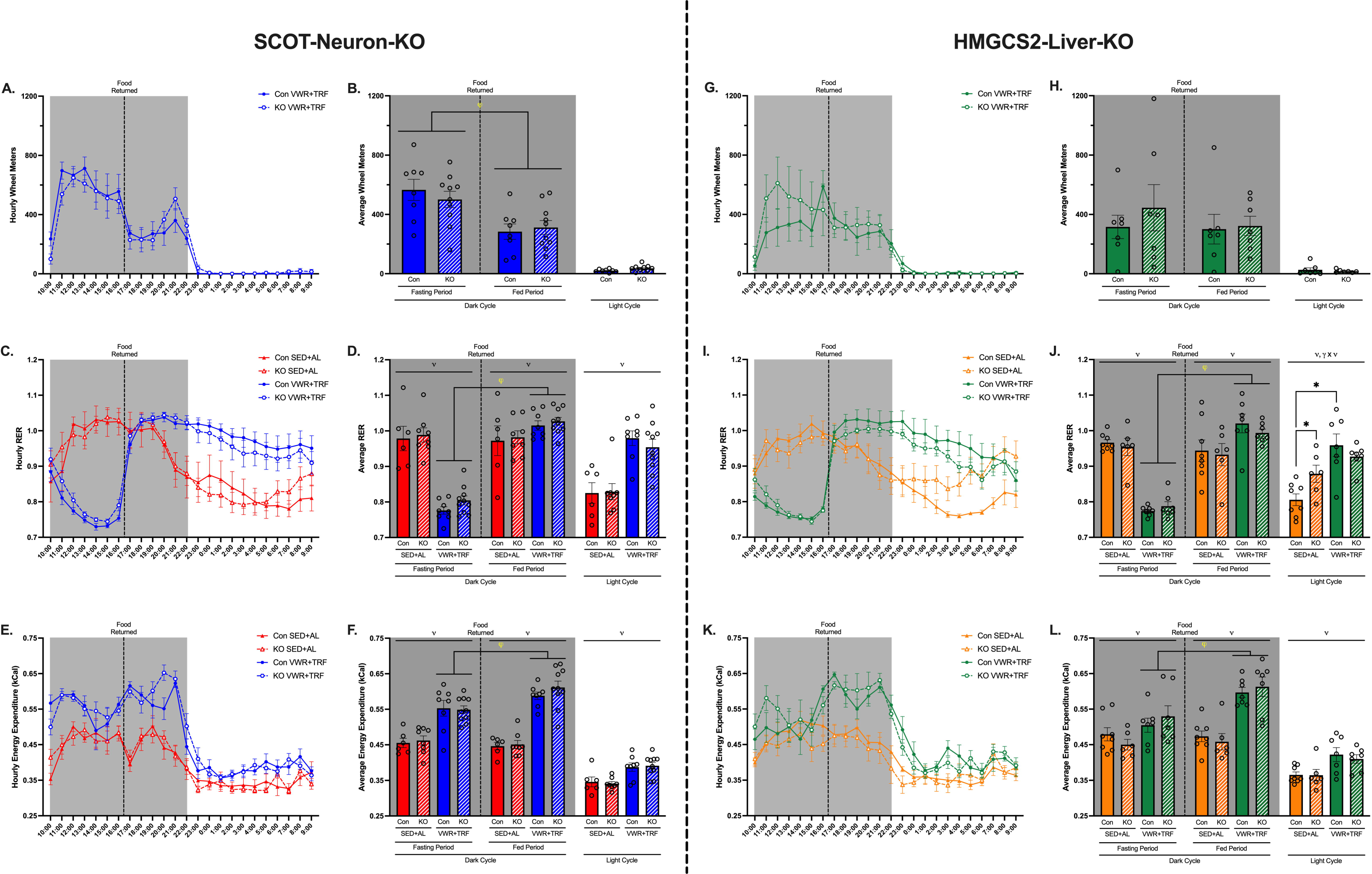
Indirect calorimetry in SCOT-Neuron-KO and HMGCS2-Liver-KO female mice. (A-F) SCOT-Neuron-KO and Con mice. (G-L) HMGCS2-Liver-KO and Con mice. (A/G) Hourly and (B/H) average wheel meters. (C/I) Hourly and (D/J) average respiratory exchange ratio (RER). (E/K) Hourly and (F/L) average energy expenditure. Data are represented as means ± SE. *n* = 6-10 per group. ϖ represents main effect of VWR+TRF following Two-way ANOVA. χπ represents main effect of fasting following Two-way ANOVA. γ x ϖ indicates significant interaction between genotype and VWR+TRF following Two-way ANOVA, with * denoting significant LSD-post hoc test.

**Figure 3.**
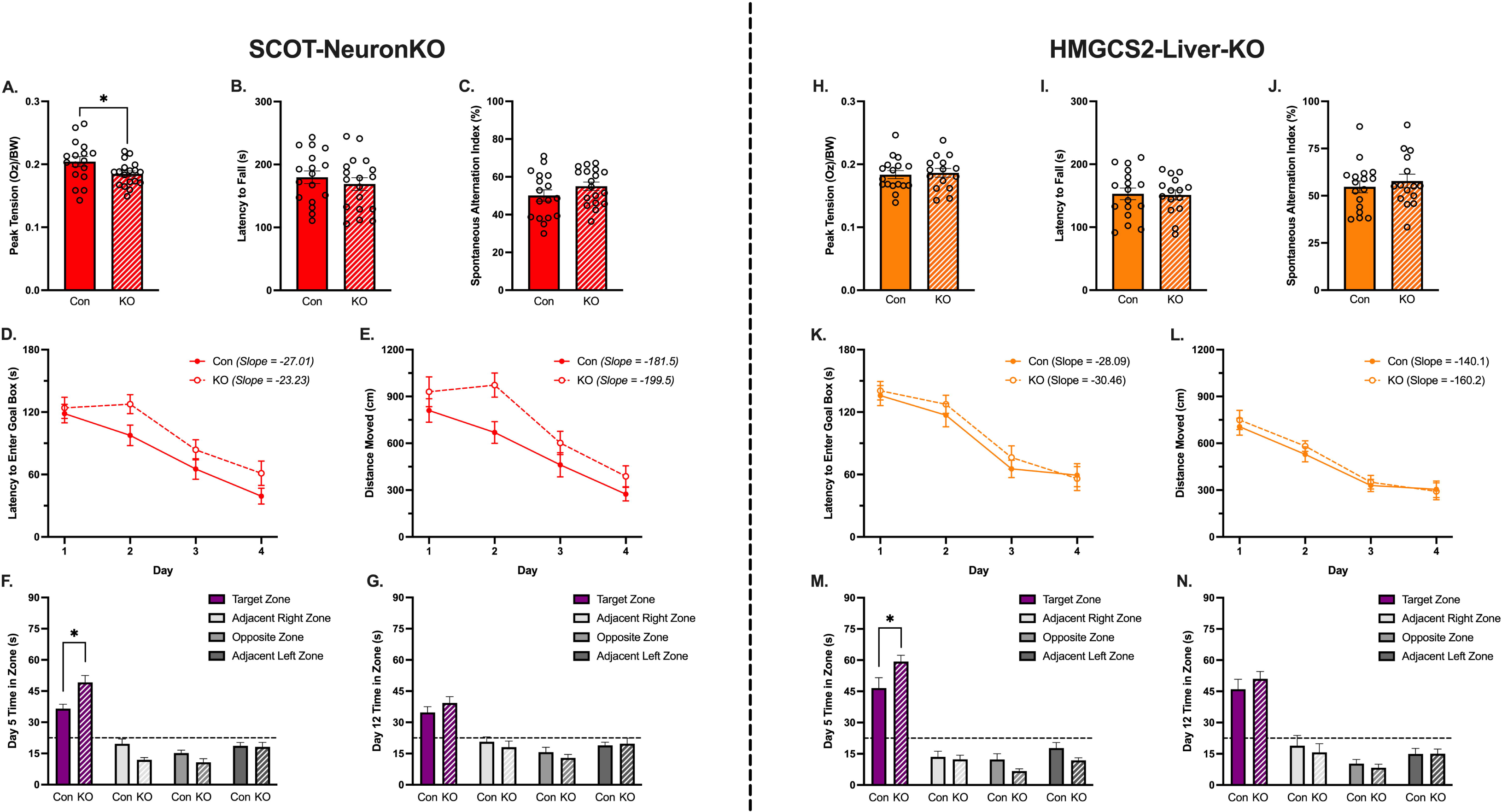
**Baseline differences in motor and cognitive behavior testing in SCOT-Neuron-KO and HMGCS2-Liver-KO female mice.** (A-G) SCOT-Neuron-KO and Con mice. (H-N) HMGCS2-Liver-KO and Con mice. (A/H) Normalized forelimb grip strength, (B/I) rotarod performance, and (C/J) spontaneous alternation index calculated from modified Y-maze testing. (D-G & K-N) Barnes Maze Testing. Spatial acquisition/learning phase, Day 1 through 4, quantified by (D/K) average daily latency to enter the goal box and (E/L) average daily distance moved. Short- and long-term reference memory probes, time spent in each quadrant on (F/M) Day 5 and (G/N) Day 12. Data are represented as means ± SE. *n* = 15-18 per group. * indicates a significant difference via unpaired t-test (p < 0.05).

Similar to SCOT-Neuron-KO mice, there were no differences in running distance between female HMGCS2-Liver-KO & Con mice in the VWR+TRF groups (**Fig. 2G-H**). Importantly, both genotypes ran equivalently during the fasting period of the dark cycle (**Fig. 2G-H**). As expected, VWR+TRF decreased RER (**Fig. 2I-J**, main effect of VWR+TRF p<0.0001) and increased EE (**Fig. 2K-L**, main effect of VWR+TRF p<0.05) during the fasting period of the dark cycle in both female HMGCS2-Liver-KO & Cons compared to SED+AL. After food was returned, there were no differences between genotypes in running distance (**Fig. 2G-H**). However, following the return of food, both RER (**Fig. 2I-J**, main effect of fasting p<0.0001) and EE (**Fig. 2K-L**, main effect of fasting p<0.01) increased within VWR+TRF groups in both female HMGCS2-Liver-KO & Cons. During the fed period of the dark cycle, VWR+TRF increased RER (**Fig. 2I-J**, main effect of VWR+TRF p<0.05) and EE (**Fig. 2K-L**, main effect of VWR+TRF p<0.0001) in female HMGCS2-Liver-KO & Con mice. During the light cycle, VWR+TRF also sustained an increased RER (**Fig. 2I-J**, main effect of VWR+TRF p=0.0001) and EE (**Fig. 2K-L**, main effect of VWR+TRF p<0.01). However, RER remained significantly more elevated in female HMGCS2-Liver-KOs compared to Cons within SED+AL, and the VWR+TRF mediated increases in RER in Cons was blunted in HMGCS2-Liver-KOs (**Fig. 2I-J**, VWR+TRF and genotype interaction p<0.05). Like in SCOT-Neuron-KO mice, VWR+TRF also pairs ketogenesis with increased energy expenditure in HMGCS2-Liver-KO mice, independent of genotype effects.

### Both SCOT-Neuron-KO and HMGCS2-Liver-KO mice display improved short-term memory at baseline

Baseline behavior and physical function testing was conducted to establish whether the developmental loss of SCOT or HMGCS2 alters motor and cognitive function. Female SCOT-Neuron-KO mice exhibited minimally reduced forelimb grip strength measured with a digital force gauge (**Fig. 3A**, p<0.05), but similar motor coordination and balance compared to Con mice measured by rotarod (**Fig. 3B**). Cognitively, female SCOT-Neuron-KO & Con mice displayed analogous spatial working memory based on Y-maze performance (**Fig. 3C**) and spatial acquisition/learning rates assessed with Barnes Maze testing (**Fig. 3D-E**). However, female SCOT-Neuron-KO mice demonstrated improved short-term memory compared to Con mice measured with Barnes Mazes recall testing (**Fig. 3F**, p<0.01). This difference did not persist into long-term memory (**Fig. 3G**). Male SCOT-Neuron-KO mice had similar grip strength (**Supp. Fig. 4A**) plus motor coordination and balance (**Supp. Fig. 4B**) as Con mice, but diminished spatial working memory (**Supp.** **Fig. 4C**, p<0.05). There was also no difference with regard to the rate of spatial acquisition/learning, short-, or long-term memory between male SCOT-Neuron-KO & Con mice (**Supp. Fig. 4D-G**). Female HMGCS2-Liver-KO mice exhibited similar forelimb grip strength, motor coordination and balance, spatial working memory, and spatial acquisition/learning rate compared to Con mice (**Fig. 3H-L**). Similar to female SCOT-Neuron-KO mice, female HMGCS2-Liver-KO mice demonstrated improved short-term memory compared to Con mice (**Fig. 3M**, p<0.05), a phenotype that also did not persist into long-term memory (**Fig. 3N**).

**Figure 4.**
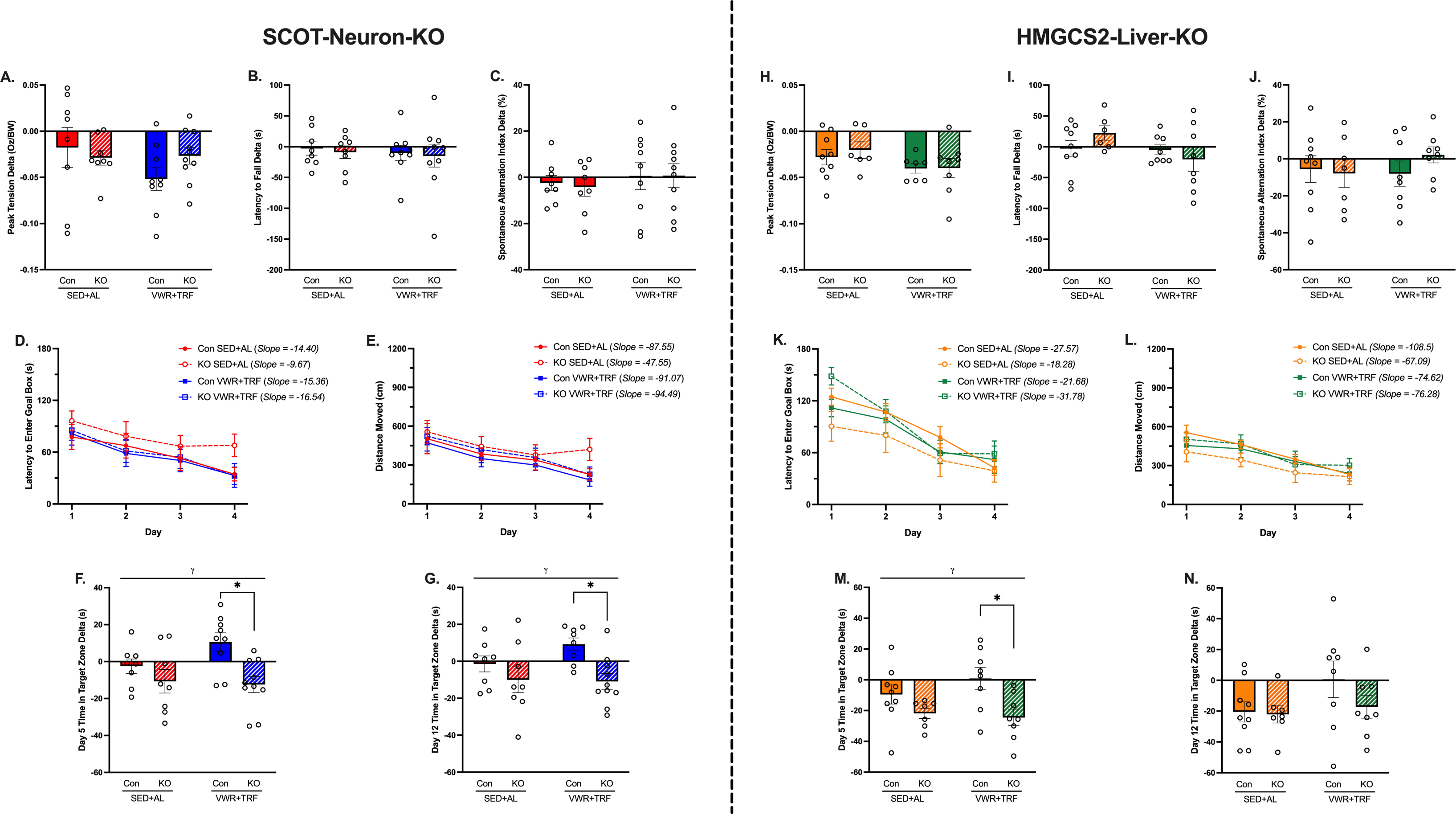
VWR+TRF reveals latent cognitive deficits in SCOT-Neuron-KO and HMGCS2-Liver-KO female mice. (A-G) SCOT-Neuron-KO and Con mice. (H-N) HMGCS2-Liver-KO and Con mice. Change from baseline in (A/H) normalized forelimb grip strength, (B/I) rotarod performance, and (C/J) spontaneous alternation index. Spatial acquisition/learning phase of endpoint Barnes Maze testing, Day 1 through 4, quantified by (D/K) average daily latency to enter the goal box and (E/L) average daily distance moved. Change from baseline in time spent in target quadrant on (F/M) Day 5 and (G/N) Day 12 during short- and long-term reference memory probes, respectively. Data are represented as means ± SE. *n* = 6-10 per group. γ represents main effect of genotype following Two-way ANOVA, with * denoting significant LSD-post hoc test.

### SCOT-Neuron-KO and HMGCS2-Liver-KO mice exhibit impaired responses to VWR+TRF in memory outcomes

We conducted the same battery of motor and cognitive behavior testing to assess whether expected decreases in performance as a result of aging were mitigated by VWR+TRF and/or impacted by impaired ketone body metabolism. No discernible differences were observed in forelimb grip strength, motor coordination and balance, or spatial working memory in female SCOT-Neuron-KO & Con mice, regardless of intervention (**Fig. 4A-C**). While we also observed no differences in changes in forelimb grip strength or motor coordination and balance in male SCOT-Neuron-KO & Con mice (**Supp. Fig. 5A-B**), male SCOT-Neuron-KO mice showed improved spatial working memory compared to Con mice, independent of intervention (**Supp. Fig. 5C**, main effect of genotype p<0.05). Female SCOT-Neuron-KO & Con mice showed no differences in spatial acquisition/learning rates regardless of intervention (**Fig. 4D-E**). However, in male SCOT-Neuron-KO & Con mice, we observed a significant interaction between genotype and VWR+TRF in the rates of spatial acquisition/learning when quantified by the latency to enter the goal-box (**Supp. Fig. 5D-E**, p<0.05). VWR+TRF led to a significantly faster rate of spatial acquisition/learning in male Con mice (**Supp. Fig. 5D-E**, p<0.01), an effect which did not occur in male SCOT-Neuron-KOs despite undergoing VWR+TRF, however this effect was largely driven by poorer performance in male Con mice in VWR+TRF on Day 1 (**Supp. Fig. 5D-E**, p<0.01). This phenotype was also not reflected in the rate of spatial acquisition/learning when quantified by the distance moved (**Supp. Fig. 5F-G**). Female SCOT-Neuron-KO mice exhibited diminished short- (**Fig. 4F**, main effect of genotype p<0.01) and long-term (**Fig. 4G**, main effect of genotype p<0.01) memory compared to Con mice. This phenotype was predominantly driven by divergent responses to VWR+TRF between female SCOT-Neuron-KO & Cons (**Fig. 4F-G**, p<0.01). Male SCOT-Neuron-KO & Con mice displayed no differences in short- or long-term memory following VWR+TRF (**Supp. Fig. 5H-I**).

**Figure 5.**
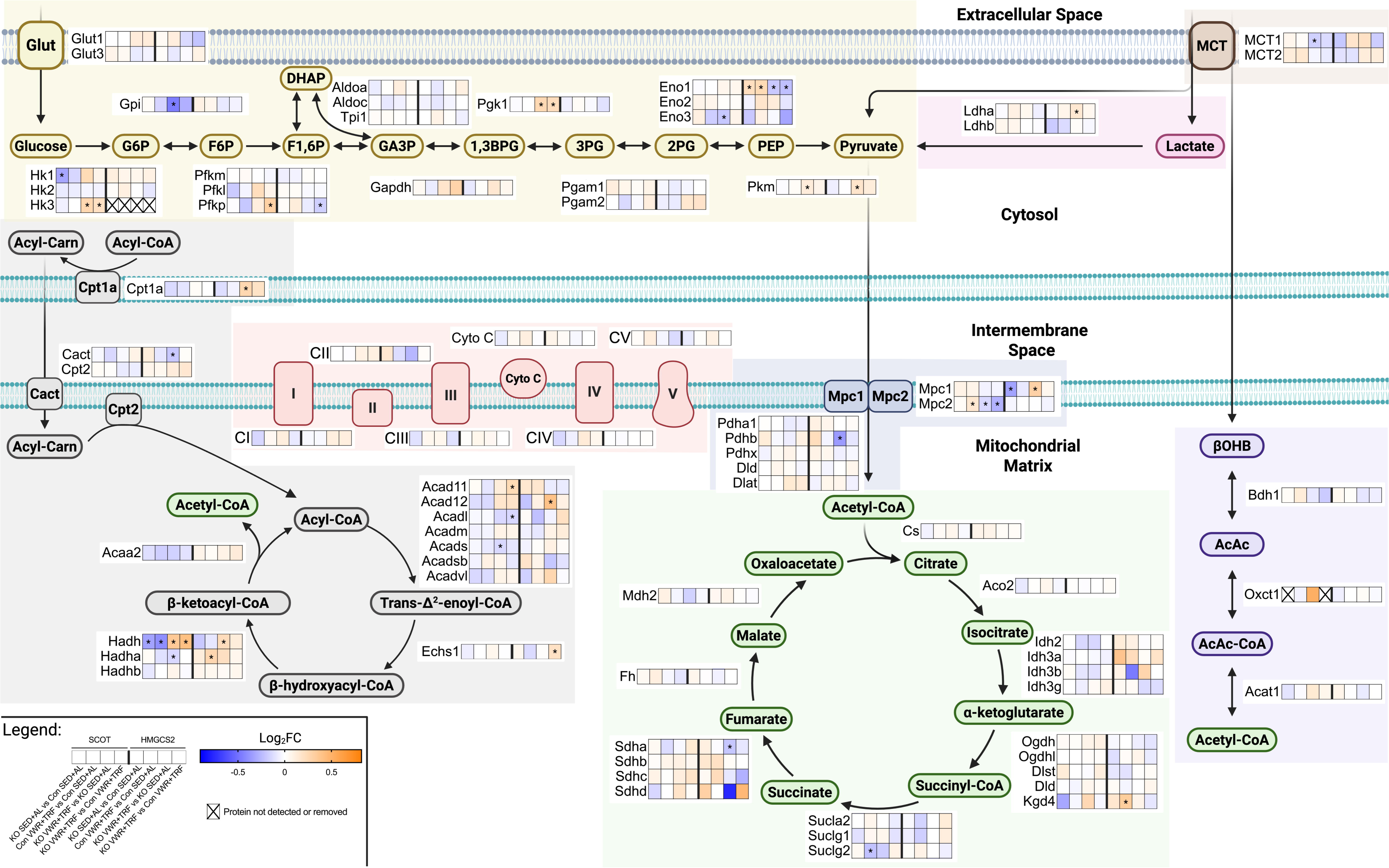
Altered hippocampal metabolic protein expression in SCOT-Neuron-KO female mice. Log_2_FC of proteins in common metabolic processes: Glucose transport, Glycolysis, Pyruvate Metabolism, Lactate Dehydrogenase, Monocarboxylate Transporters, Ketolysis, TCA cycle, Fat Transport, β-oxidation, and Oxidative Phosphorylation. * indicates significant p-value (p<0.05). BioRender was used to generate figure.

Female HMGCS2-Liver-KO & Con mice demonstrate no differences in changes in forelimb grip strength, motor balance and coordination, or spatial working memory after VWR+TRF (**Fig. 4H-J**). Female HMGCS2-Liver-KO & Con mice also display similar rates of spatial acquisition/learning at endpoint, regardless of intervention (**Fig. 4K-L**). Female HMGCS2-Liver-KO mice exhibit reductions in short-term memory compared to Con mice (**Fig. 4M**, main effect of genotype p<0.01), which is also driven by divergent responses to VWR+TRF between HMGCS2-Liver-KO and Con mice (**Fig. 4M**, p<0.01). However, this phenotype did not persist into long-term memory as it did in female SCOT-Neuron-KO & Con mice (**Fig. 4N**). Collectively, these data show that despite superior short-term memory performance at baseline, both female SCOT-Neuron-KO and HMGCS2-Liver-KO mice display divergent responses to chronic VWR+TRF compared to their respective Con mice, resulting in a more significant decline in short-term memory with aging.

### Female SCOT-Neuron-KO and HMGCS2-Liver-KO mice show uniquely altered hippocampal proteomic responses to VWR+TRF

To determine the molecular underpinnings of impaired responses to VWR+TRF in indices of short- and long-term memory in SCOT-Neuron-KO mice, we performed bulk proteomics on the right hippocampus of female SCOT-Neuron-KO & Con mice, as well as HMGCS2-Liver-KO & Con mice (n = 4-5 per group, quality control conducted by IDeA National Resource for Quantitative Proteomics removed 1 sample in the SCOT Con SED+AL group). Comparisons within SCOT-Neuron-KO vs Con and within HMGCS2-Liver-KO vs Con included: KO SED+AL vs Con SED+AL (baseline effects of KO), Con VWR+TRF vs Con SED+AL (effect of VWR+TRF within Con mice), KO VWR+TRF vs KO SED+AL (effect of VWR+TRF within KO), and KO VWR+TRF vs Con VWR+TRF (effect of KO within VWR+TRF). We first surveyed proteins of interest linked to glucose, fat, and ketone body metabolism. Changes at the individual protein level for metabolic processes (glucose transport, glycolysis, lactate dehydrogenase, pyruvate metabolism, monocarboxylate transporters, ketolysis, TCA cycle, fat transport, β-oxidation, and oxidative phosphorylation) are shown in **Figure 5**. Within SCOT-Neuron-KO vs Con mice, SCOT (*Oxct1*) expression comparisons between Con and KO groups were removed as not to drive the scale of Log_2_FC for all other comparisons (SCOT-Neuron-KO SED+AL vs Con SED+AL, Log_2_FC = -1.91 and SCOT-Neuron-KO VWR+TRF vs Con VWR+TRF, Log_2_FC = -1.35). We then used the normalized summation of proteins in each metabolic process depicted in **Figure 5** to evaluate changes across all groups within SCOT-Neuron-KO vs Con mice and within HMGCS2-Liver-KO vs Con mice. In SCOT-Neuron-KO vs Con mice, there were no differences in total protein content between our groups (**Fig. 6A**). While there were no differences in glucose transport (**Fig. 6B**), SCOT-Neuron-KO mice demonstrated an overall increased expression of proteins involved in glycolysis (**Fig. 6C**, main effect of genotype p<0.05). This effect was predominantly driven by VWR+TRF upregulation of glycolytic proteins in SCOT-Neuron-KO but not Con mice (**Fig. 6C**, p<0.01). VWR+TRF showed a modest, non-statistically significant increase (p=0.09) in monocarboxylate transporter expression in Con mice, but not in SCOT-Neuron-KO mice. Under SED+AL conditions, SCOT-Neuron-KO mice showed a modest, non-statistically significant increase (p=0.06) in monocarboxylate transporter expression compared to Con mice, although neither of these pairwise comparisons reached significance (**Fig. 6D**, genotype and VWR+TRF interaction p<0.05). As expected, a marked decrease in the expression of ketolysis proteins was observed in the hippocampi of SCOT-Neuron KOs compared to Con mice (**Fig. 6E**, main effect of genotype p<0.0001), without an effect of VWR+TRF in Con mice. Interestingly, VWR+TRF caused a modest increase in global protein expression in the ketolysis pathway in SCOT-Neuron-KO mice, suggesting possible compensatory increases in ketolysis in glial cell populations with VWR+TRF (**Fig. 6E**, main effect of VWR+TRF p=0.0502). VWR+TRF also tended to increase the expression of proteins in β-oxidation, but only in SCOT-Neuron-KO mice, and this pairwise comparison did not reach significance (**Fig. 6F**, genotype and VWR+TRF interaction p<0.05). There were no differences in the other metabolic processes that were evaluated in SCOT-Neuron-KO & Con mice (**Supp. Fig. 6A-J**). Within HMGCS2-Liver-KO & Cons, we observed no differences in total protein content (**Fig. 6G**) or with any of the metabolic processes evaluated, regardless of genotype or intervention (**Fig. 6H-L** and **Supp. Fig 6K-T**).

**Figure 6.**
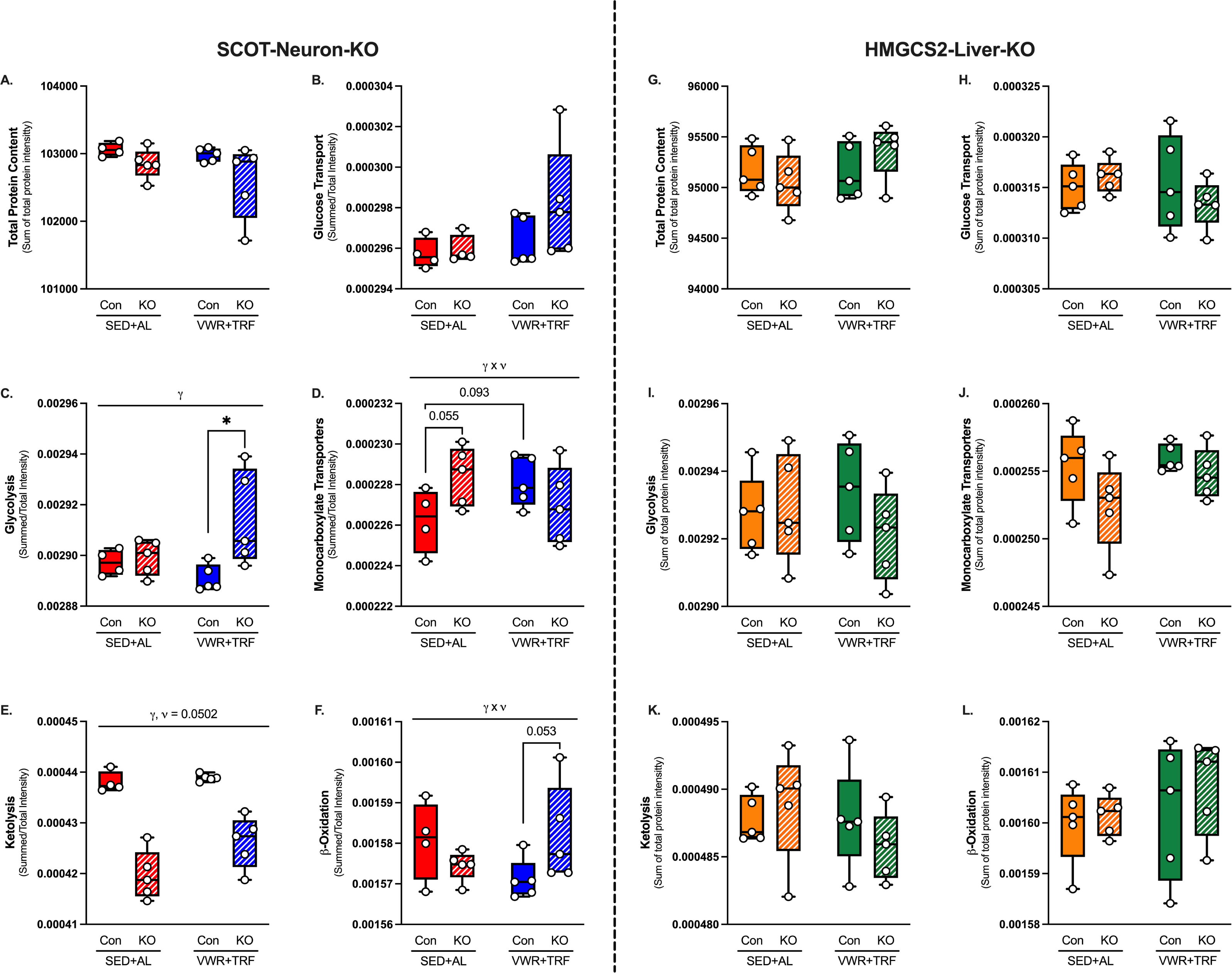
Differential hippocampal proteomic signatures of metabolic adaptations in SCOT-Neuron-KO and HMGCS2-Liver-KO female mice. (A) Total VSN normalized exclusive intensity of all proteins within each sample, i.e., total protein content per sample. Summed VSN normalized exclusive intensities of proteins in each respective metabolic process, identified in Figure 5, normalized to total protein content per sample. (B) Glucose Transport, (C) Glycolysis, (D) Monocarboxylate Transporters, (E) Pyruvate Metabolism, (F) Lactate Dehydrogenase, (G) Ketolysis, and (H) β-oxidation. Data are represented as means ± SE. *n* = 4-5 per group. ϖ and γ represent main effects of VWR+TRF and genotype, respectively, following Two-way ANOVA. γ x ϖ indicates significant interaction between genotype and VWR+TRF following Two-way ANOVA. * denotes significant LSD-post hoc test.

### VWR+TRF upregulates synaptic machinery in Con mice, but not SCOT-Neuron-KO mice

To perform an unbiased interrogation of the hippocampal proteome in response to VWR+TRF and the loss of SCOT in neurons, we performed a pathway analysis using the Log_2_FC and p-values of the differentially expressed proteins with IPA. The top 10 up- and down-regulated pathways identified in each comparison, ranked by Z-score, are displayed in **Supplemental Figure 7A-D**. We focused on differentially regulated pathways within the SCOT-Neuron-KO VWR+TRF vs Con VWR+TRF comparison, given that our behavior phenotypes were predominantly driven by a divergence between these two groups. Significant downregulations of (1) neurexins and neuroligins, and (2) assembly and cell surface presentation of NMDA receptors pathways represented core alterations signifying perturbations of synaptic maturation. Therefore, heat maps were generated displaying the individual Log_2_FC values of proteins that IPA identified in these 2 pathways for all comparisons within SCOT-Neuron-KO & Con mice (**Fig. 7A & F**). We divided each pathway’s proteins into clusters based on function and used the normalized summation of each functional cluster to evaluate changes across all groups within SCOT-Neuron-KO & Con mice. Within the neurexins and neuroligins pathway, there were no differences in synaptic scaffolding and adaptor proteins or synaptic receptor expression (**Fig. 7B-C**). However, VWR+TRF increased the expression of synaptic cell adhesion molecules within Con mice, an effect that did not occur in SCOT-Neuron-KO mice (**Fig. 7D**, genotype and VWR+TRF interaction p<0.05). No differences were observed in proteins involved in vesicle trafficking and neurotransmitter release (**Fig. 7E**). Within the assembly & cell surface presentation of NMDA receptors pathway, VWR+TRF induced an increase in synaptic scaffolding and adaptor proteins in Con mice, but not in SCOT-Neuron-KO mice (**Fig. 7G**, genotype and VWR+TRF interaction p<0.01). No differences in synaptic cytoskeleton protein expression were observed between groups (**Fig. 7H**), but VWR+TRF tended to increase the expression of synaptic receptors in Con mice, but not SCOT-Neuron-KOs, however this did not reach significance (**Fig. I**, genotype and VWR+TRF interaction p=0.094). No differences in the expression of synaptic kinases between groups were observed (**Fig. 7J**). These data reveal potential molecular mechanisms by which neuronal ketone utilization may mediate the maintenance of synaptic plasticity with aging in response to chronic VWR+TRF.

**Figure 7.**
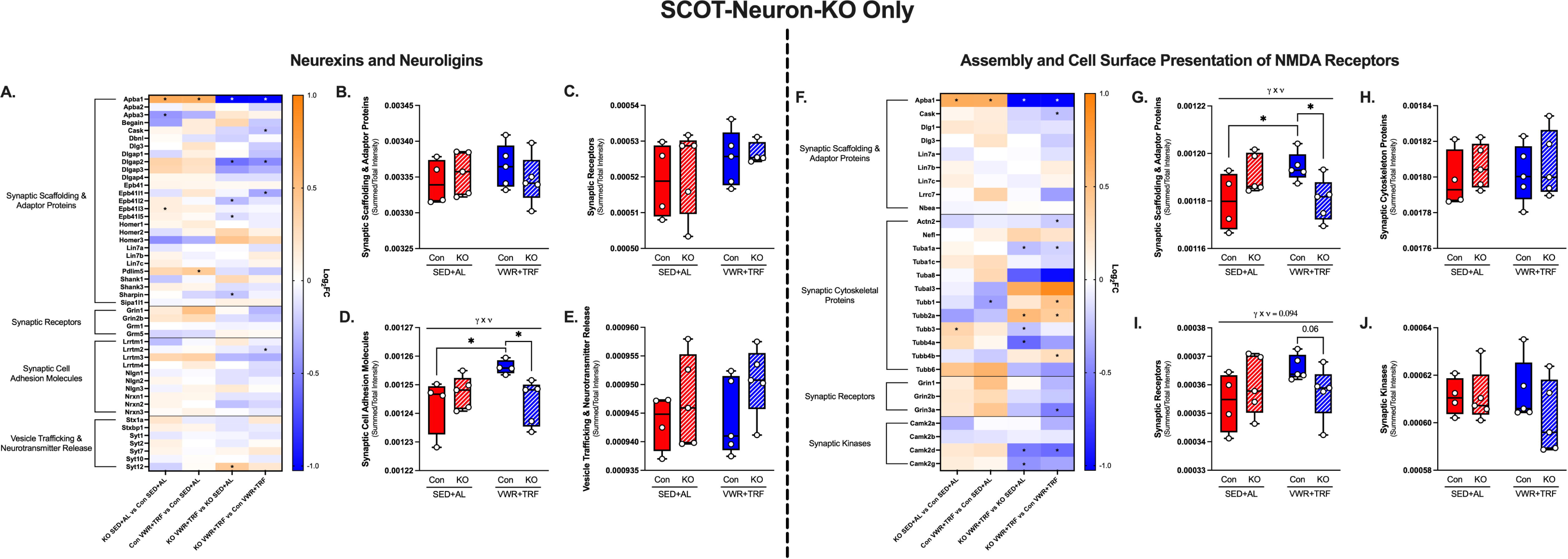
Altered hippocampal proteomic signatures of synaptic machinery in SCOT-Neuron-KO female mice. (A) Heatmaps for the Log_2_FC of individual proteins in the IPA pathway called Neurexin and Neuroligins, which was significantly downregulated in the SCOT-Neuron-KO VWR+TRF vs Con VWR+TRF comparison, i.e., up with VWR+TRF in Con. Proteins within the pathway were separated based on function. (B) Synaptic Scaffolding and Adaptor proteins, (C) Synaptic Receptors, (D) Synaptic Cell Adhesion Molecules, and (E) Vesicle Trafficking & Neurotransmitter Release. (F) Heatmaps for the Log_2_FC of individual proteins in the IPA pathway called Assembly and Cell Surface Presentation of NMDA Receptors, which was significantly downregulated in the SCOT-Neuron-KO VWR+TRF vs Con VWR+TRF comparison, i.e., up with VWR+TRF in Con. Proteins within the pathway were separated based on function. (G) Synaptic Scaffolding & Adaptor Proteins, (H) Synaptic Cytoskeleton Proteins, (I) Synaptic Receptors, (J) Synaptic Kinases. * within heatmaps signifies significant p-value (p<0.05). Data are represented as means ± SE. *n* = 4-5 per group. γ x ϖ indicates significant interaction between genotype and VWR+TRF following Two-way ANOVA, with * denoting significant LSD-post hoc test.

### HMGCS2-Liver-KO globally increases the expression of hippocampal proteins that mediate classical ketogenesis

We then performed an unbiased interrogation of the hippocampal proteome in response to VWR+TRF and the loss of HMGCS2 in the liver. The top 10 up- and down-regulated pathways identified in each comparison, ranked by Z-score, are displayed in **Supplemental Figure 7E-H**. However, no significant pathways representing perturbations in synaptic function were observed. Taken together with the lack of observed differences in the hippocampal metabolic processes we previously evaluated (**Fig. 6H-L** and **Supp. Fig 6K-T**), we sought to determine whether compensatory ketogenesis was occurring in the hippocampus of HMGCS2-Liver-KO mice. Changes at the individual protein level for classical ketogenesis are shown in **Figure 8A**. We then used the normalized summation of the proteins shown to evaluate changes across all groups within HMGCS2-Liver-KO & Con mice. Strikingly, HMGCS2-Liver-KO mice showed increased expression of classical ketogenesis proteins in the hippocampus (**Fig. 8B**, main effect of genotype p<0.01), an effect that was only observed in the SED+AL condition (**Fig. 8B** p<0.01). Overall, this finding suggests that there is a glial compensatory increase in ketogenesis in the hippocampus of liver-specific HMGCS2 KO mice.

**Figure 8.**
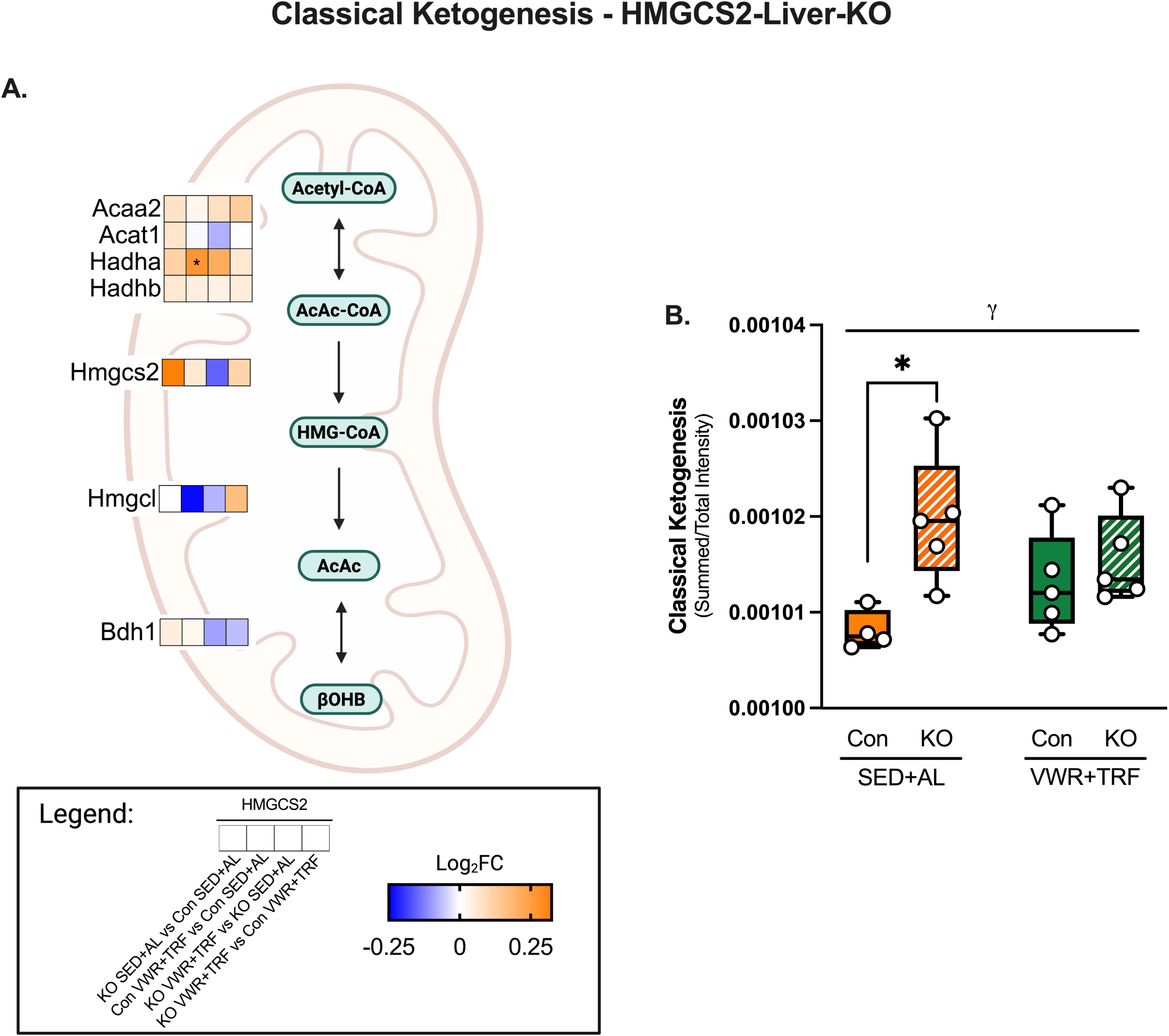
Induction of canonical ketogenesis in the hippocampus of HMGCS2-Liver-KO female mice. (A) Log_2_FC of proteins in the classical ketogenesis pathway. * indicates significant p-value (p<0.05). (B) Summed VSN normalized exclusive intensities of proteins shown in (A) normalized to total protein content per sample. Data are represented as means ± SE. *n* = 4-5 per group. γ represents a main effect of genotype following Two-way ANOVA. * denotes significant LSD-post hoc test.BioRender was used to generate figure graphic.

## Discussion

In this study, we sought to determine whether ketone bodies are molecular mediators of exercise-induced brain health benefits. We selectively targeted neuronal ketone body utilization (SCOT) and hepatic ketone body production (HMGCS2) to determine their respective roles in the responses to chronic VWR+TRF and the modulation of whole body metabolism and cognitive function in middle-aged mice. Our data are consistent with the following: (1) VWR+TRF shifts systemic metabolism toward fat-derived sources, with increased energy expenditure, stimulating robust ketone body turnover; (2) female SCOT-Neuron-KO mice show relative short- and long-term memory impairments compared to Con mice in response to VWR+TRF; (3) memory phenotypes were paired with failed upregulation of key facilitators of synaptic function within the neurexin and neuroligin pathway, and within the assembly and presentation of NMDA receptors pathway in the hippocampi of female SCOT-Neuron-KO mice. We also observed shifts in the global expression of metabolic processes as a result of VWR+TRF in female SCOT-Neuron-KO mice. In HMGCS2-Liver-KO mice, we observed diminished short-term, but not-long term memory following VWR+TRF compared to Con mice. Unlike SCOT-Neuron-KO mice, this phenotype was not matched with significant changes in indicators of synaptic function in the hippocampus via pathway enrichment analysis. However, a global increase in the expression of ketogenic enzymes in the hippocampus of HMGCS2-Liver-KO mice was observed, and when paired with our memory phenotype, suggests a possible compensatory local production of ketone bodies that should be explored in future studies.

Under normal physiological conditions, humans undergo regular circadian oscillations in ketone body concentrations between roughly 50 and 250 μM^18–20^. These oscillations are predominantly driven by the fed and fasted states associated with our circadian rhythm because the rate of ketogenesis mirrors the rate of adipose tissue lipolysis and hepatic mitochondrial β-oxidation^18–20^. This phenomenon was observed in both female and male SCOT-Neuron-KO and Con mice undergoing SED+AL. During the dark cycle as the mice are active and eating, carbohydrate-derived sources largely drive energy production, reflected in RER values at roughly 1. As the mice transitioned to the light cycle and accompanied inactivity, RER declined to about 0.8, indicating a shift toward more fat-derived energy sources. While Con mice in the SED+AL state showed this phenomenon, HMGCS2-Liver-KOs exhibited a blunted oscillation, despite similar caloric intake. This may result from compensatory increases in hepatic glycogen stores and hepatic glycogenolysis, as previously observed in an HMGCS2 knockdown model^50^. On the other hand, VWR+TRF reversed the timing of these circadian oscillations for all mice so that ketogenesis occurred during the dark rather than the light cycle. VWR+TRF effectively altered the timing of the natural circadian oscillation, pairing ketogenesis with increased energy expenditure and thereby stimulating robust ketone body turnover, unlike the oscillation in SED+AL groups^17^. Importantly, mice in VWR+TRF did have access to food for the last 5.5 hours of the dark cycle, providing partial alignment with their normal time of food consumption. VWR+TRF was also associated with modestly elevated RER during the light cycle, although with a steady downward trend. This could be a combined result of eating later in the dark cycle and exercise-induced adaptations in liver glycogen stores, allowing the VWR+TRF groups to rely more on carbohydrate-derived sources during the light cycle than SED+AL mice^51^.

Interestingly, female SCOT-Neuron-KO mice were protected from the gain in fat mass, albeit a difference that reflects less than 5% total body mass, that Con mice exhibited following VWR+TRF, despite similar energy expenditure as measured by indirect calorimetry and caloric intake throughout the study. This phenotype could largely be driven by weight loss, likely a result of behavior testing stressors, observed from week 17 to week 18 in SCOT-Neuron-KO mice in VWR+TRF. Alternatively, enhanced glucose sparing by tissues other than the brain in SCOT-Neuron-KO mice could mediate this phenotype. Previous evidence has established that neonatal neuronal SCOT-Neuron-KO mice exhibit euglycemia and hyperketonemia following 24- and 48-hour fasts and develop increased cerebral glycolysis and glucose oxidation^24^. Here we show that VWR+TRF in adult female SCOT-Neuron-KO mice increases the overall hippocampal abundance of glycolytic proteins compared to Con mice. Taken together, in female SCOT-Neuron-KO mice undergoing VWR+TRF, neurons likely compensate for the lack of terminal ketone oxidative capacity by increasing glucose utilization. This may require enhanced glucose sparing by other metabolic tissues, including compensatory increases in ketone body utilization in depots like cardiac and skeletal muscle, as suggested in a previous report^24^. Skeletal muscle may also spare glucose by increasing reliance on fat oxidation. The selective deletion of mitochondrial pyruvate carrier in skeletal muscle increases fatty acid oxidation, blunting the expansion of adiposity^52^.

Furthermore, our proteomics data suggest that VWR+TRF in SCOT-Neuron-KO mice also tends to globally upregulate the expression of proteins in β-oxidation within the hippocampus. While bulk proteomics cannot distinguish among cell types, because neurons are relatively unable to perform β-oxidation, our data suggest that other cell types in the brain may also be sparing glucose for neurons^53^. These adaptations are putatively linked to our observed body composition phenotype, as adipose tissue lipolysis may be enhanced to meet energetic demand while sparing glucose, especially in the background of VWR+TRF. However, our data would suggest that these adaptations are sex specific, given that male SCOT-Neuron-KO mice undergoing VWR+TRF do not display the same changes in body composition. Rather, male SCOT-Neuron-KO mice in SED+AL do.

We also evaluated how the developmental loss of neuronal SCOT or hepatic HMGCS2, and therefore targeted perturbations in ketone body metabolism, may impact motor and cognitive function prior to our intervention. Only female SCOT-Neuron-KO mice presented with minimally diminished forelimb grip strength compared to their Con counterparts, despite similar body composition. This phenotype did not persist in motor coordination and balance or average daily running distance, suggesting the differences in grip strength did not drive major differences in gross function. Female SCOT-Neuron-KO and HMGCS2-Liver-KO mice both exhibited improved short- but not long-term memory compared to their Con counterparts at baseline, despite learning the Barnes Maze task at the same rate and reaching similar levels of proficiency on Day 4. While HMGCS2-Liver-KO mice maintain euglycemia when fasted, no current evidence explores substrate utilization in the brain^32^. However, previous evidence has shown increased glucose oxidation in the brains of neonatal neuronal SCOT-Neuron-KO mice^24^. It is possible that this adaptation yields positive effects on short-term memory processes compared to Con mice, in a sex-dependent manner, as male SCOT-Neuron-KO mice displayed reduced spatial working memory compared to Cons, but similar rates of learning, short-, and long-term memory.

Following intervention, we found diminished short- and long-term memory in female SCOT-Neuron-KOs compared to Con mice, regardless of VWR+TRF. This also occurred despite similar learning rates and Day 4 performance at endpoint Barnes Maze. Although, here, we are unable to statistically show that VWR+TRF significantly improved short- and long-term memory in Cons, the divergent responses between SCOT-Neuron-KO & Cons to VWR+TRF primarily drive this phenotype. Future studies should explore the effect of chronic VWR+TRF in elderly mice, where we would expect a more pronounced phenotype to emerge in Cons. Nonetheless, these data suggest that neuronal terminal ketone body oxidation is vital in maintaining memory function during physiological aging and shows important promise as therapeutic “exerkine”. To reveal the molecular mechanisms driving these cognitive phenotypes, we performed bulk proteomics on the right hippocampus. Pathway analysis revealed that VWR+TRF failed to upregulate neurexin and neuroligin and assembly and cell surface presentation of NMDA receptors pathways in SCOT-Neuron-KO mice like it did in Con mice. After dividing the proteins within the neurexin and neuroligin pathway into functional clusters, we found that VWR+TRF in Cons globally upregulated synaptic cell adhesion molecules, namely, leucine-rich repeated transmembrane (Lrrtm) proteins 1-4, neuroligins (Nlgn) 1-3, and neurexins (Nrxn) 1-3. These synaptic cell adhesion molecules have been shown to interact with one another, especially Lrrtm2 and Nrxn1, and are linked to synaptic formation, maturation, plasticity, and function^54–60^. The knockdown of Lrrtm1 in mice mediated a reduction in both hippocampal volume and synaptic density as well as elicited short-term memory impairments, while the double knockdown of Lrrtm1 and 2 disrupted long-term potentiation (LTP) in both neonatal and mature hippocampal neurons^54, 55^. Lrrtm3 has also been shown to play a direct role in synaptic density and strength in the dentate gyrus of the hippocampus^57^. Knockout models of Nlgns and Nrxns have also shown detriments to synaptic transmission^60^.

Nlgn1 has even been explored in the context of neurodegeneration, and its expression is found to be diminished in the hippocampus of humans with Alzheimer’s disease compared to healthy controls^56^. VWR+TRF in Cons also globally upregulated synaptic scaffolding and adaptor proteins within the assembly and cell surface presentation of NMDA receptors pathway. Albeit these proteins are not as extensively explored in animal models, they generally function to cluster and anchor NMDA receptors in the synapse, and one study in drosophila demonstrated that defects in discs large homolog (dlg) hindered short-term memory^61^. While our study does not directly capture *in vivo* or *ex vivo* measurements of synaptic function/transmission or LTP, these proteomic findings in combination with our functional behavior outcomes provide a possible mechanism by which ketone bodies mediate cognitive benefit as exerkines and warrant exploration in future studies.

We expected a more robust phenotype in the female HMGCS2-Liver-KOs compared to Con mice due to a systemic deficit in circulating ketone bodies, however, we only observed diminished short-term memory, regardless of VWR+TRF. This also occurred despite similar learning rates and Day 4 performance at endpoint Barnes Maze.

Furthermore, we failed to identify significant enrichment of pathways related to synapses or synaptic function. We also did not observe prominent metabolic adaptations like those in the SCOT-Neuron-KO mice. A more prominent memory phenotype and potential molecular mechanisms in female SCOT-Neuron-KOs compared to HMGCS2-Liver-KOs could provide novel *in vivo* evidence of a possible physiological role of local ketone body synthesis in the brain that compensates for impaired hepatic ketogenesis. In support of this hypothesis, our proteomics data showed a global increase in hippocampal expression of proteins involved in the classical ketogenic pathway in liver specific HMGCS2-Liver-KO mice, driven primarily by differences in the SED+AL groups. This could indicate glial compensation for a lack of hepatic ketogenesis. Indeed, an astrocyte-neuron ketone body shuttle has been postulated and is the most explored possibility^62^. β-oxidation within the central nervous system largely occurs in astrocyte populations, and is mostly excluded from neurons, although this latter position has been recently challenged^53, 63, 64^. Octanoate, a medium chain fatty acid, has been shown to be readily oxidized in the brain *in vivo*, which is in part attributed to its transport not being limited by carnitine-acylcarnitine translocase^65^. Astrocytes have been shown to produce ketone bodies *in vitro*, especially when provided with octanoate, and have therefore been proposed to provide a localized source of ketones in the brain^66–69^. There are, however, several notable caveats to this proposed shuttle; (1) robust evidence of astrocytic ketone body production is relatively limited to *in vitro* cultures, albeit some *in vivo* evidence does exist in the ventromedial hypothalamus of rats^66–72^. (2) Astrocytes, unlike hepatocytes, express SCOT and can therefore readily oxidize ketone bodies^66–72^. (3) The astrocytic ketogenesis pathway remains largely unknown. Prior to our observations, it was not clear whether astrocytes express HMGCS2 and therefore perform ketogenesis via the classical pathway^66–72^. An alternative ketogenic pathway is feasible by reverse flux through SCOT, which catalyzes the equilibrium reaction between AcAc and AcAc-CoA^18, 19^. In fact, the hydrolysis of AcAc-CoA into AcAc, and therefore the formation of ketones, is thermodynamically favored^18, 19^. It is the combination of energetic demand and AcAc mass action that drives the formation of AcAc-CoA and subsequent conversion to Acetyl-CoA in extrahepatic tissues^18, 19^. It is feasible that astrocyte ketogenesis is an adaptive response that occurs in HMGCS2-Liver-KO mice given that they lack the physiological AcAc mass action. While our proteomics detected HMGCS2, we are unable to connect expression to specific cell types. Taken together, astrocytes or other glia may directly contribute ketone bodies to the local *in vivo* milieu under physiological conditions like VWR+TRF or ketogenic diets. Flux analysis using stable isotope tracing could be used in future studies to better tease out cell-specific contributions in the brain.

Here, we determined the role of ketone bodies as molecular mediators of exercise-induced brain health benefits in chronic VWR+TRF with aging. Through targeting neuronal ketone body utilization, we show impaired responses to VWR+TRF in functional outputs of memory and the expression of facilitators of synaptic function. We also provide potential *in vivo* evidence of cerebral ketogenesis, given the lack of robust memory and synaptic expressional profile differences, combined with an upregulation in classical ketogenesis machinery within the hippocampus, after targeting hepatic ketone body production.

## Supplementary Figure Legends

**Figure S1. Body mass and composition in SCTO-Neuron-KO male mice.** (A) Weekly body mass. (B) Total food consumed over the duration of the study. (C) Average daily running distance. Change from baseline in (D) body mass, (E) fat free mass, (F) fat mass, and (G) percent fat mass, calculated by dividing fat mass by body mass. Data are represented as means ± SE. *n* = 6-9 per group. ϖ and γ represent main effects of VWR+TRF and genotype, respectively, following Two-way ANOVA.

**Figure S2. Indirect calorimetry in SCOT-Neuron-KO male mice.** (A) Hourly and (B) average wheel meters. (C) Hourly and (D) average RER. (E) Hourly and (F) average energy expenditure. Data are represented as means ± SE. *n* = 6-8 per group. ϖ, γ, and χπ represent main effects of VWR+TRF, genotype, and fasting, respectively, following Two-way ANOVA.

**Figure S3. Static blood D-βOHB measurements.** (A) Female and (B) male SCOT-Neuron-KO and Con mouse blood ketone concentration following a 24 hour fast in VWR+TRF groups. Data are represented as means ± SE. *n* = 5-9 per group.

**Figure S4. Baseline behavior testing in SCOT-Neuron-KO male mice.** (A) Normalized forelimb grip strength, (B) rotarod performance, and (C) spontaneous alternation index calculated from modified Y-maze testing. (D-G) Barnes Maze Testing. Spatial acquisition/learning phase, Day 1 through 4, quantified by (D) average daily latency to enter the goal box and (E) average daily distance moved. Short- and long-term reference memory probes, time spent in each quadrant on (F) Day 5 and (G) Day 12. Data are represented as means ± SE. *n* = 14-15 per group. * indicates a significant difference via unpaired t-test (p < 0.05).

**Figure S5. VWR+TRF reveals changes in motor and latent cognitive deficits in SCOT-Neuron-KO male mice.** Change from baseline in (A) normalized forelimb grip strength, (B) rotarod performance, and (C) spontaneous alternation index. Spatial acquisition/learning phase of endpoint Barnes Maze testing, Day 1 through 4, quantified by (D) average daily latency to enter the goal box (E) slope of average daily latency to enter goal box, (F) average daily distance moved, and (G) slope of average daily distance moved. Change from baseline in time spent in target quadrant on (H) Day 5 and (I) Day 12 during short- and long-term reference memory probes, respectively. Data are represented as means ± SE. *n* = 6-9 per group. γ represents main effect of genotype following Two-way ANOVA. γ x ϖ indicates significant interaction between genotype and VWR+TRF following Two-way ANOVA, with * denoting significant LSD-post hoc test.

**Figure S6. Normal hippocampal proteomic signatures of metabolic adaptions in SCOT-Neuron-KO and HMGCS2-Liver-KO female mice.** (A-H) SCOT-Neuron-KO and Con. Summed VSN normalized exclusive intensities of proteins in each respective metabolic process, identified in Figure 5, normalized to total protein content per sample. (A) TCA Cycle, (B) Fat Transport, (C) Complex I, (D) Complex II, (E) Complex III, (F) Cytochrome C, (G) Complex IV, and (H) Complex V. (I-W) HMGCS2-Liver-KO and Con. (I) Total VSN normalized exclusive intensity of all proteins within each sample, i.e., total protein content per sample. Summed VSN normalized exclusive intensities of proteins in each respective metabolic process normalized to total protein content per sample. (J) Glucose Transport, (K) Glycolysis, (L) Monocarboxylate Transporters, (M) Pyruvate Metabolism, (N) Lactate Dehydrogenase, (O) Ketolysis, (P) TCA Cycle, (Q) Fat Transport, (R) Complex I, (S) Complex II, (T) Complex III, (U) Cytochrome C, (V) Complex IV, and (W) Complex V. Data are represented as means ± SE. *n* = 4-5 per group.

**Figure S7. Unbiased pathway analysis of proteomic signatures in SCOT-Neuron-KO and HMGCS2-Liver-KO female mice.** (A-D) SCOT-Neuron-KO and Con, (E-H) HMGCS2-Liver-KO and Con. The 10 most up- and down-regulated pathways based on Z-score for each given comparison. (A) SCOT-Neuron-KO SED+AL vs Con SED+AL, (B) Con VWR+TRF vs CON SED+AL, (C) SCOT-Neuron-KO VWR+TRF vs SCOT-Neuron-KO SED+AL, (D) SCOT-Neuron-KO VWR+TRF vs Con VWR+TRF, (E) HMGCS2-Liver-KO SED+AL vs Con SED+AL, (F) Con VWR+TRF vs Con SED+AL, (G) HMGCS2-Liver-KO VWR+TRF vs HMGCS2-Liver-KO VWR+TRF, and (H) HMGCS2-Liver-KO VWR+TRF vs Con VWR+TRF. Pathways with a Z-score greater than or equal to an absolute value of 2 and with a -log_10_p-value greater than 1.3 were considered significant.

**Table S1.**
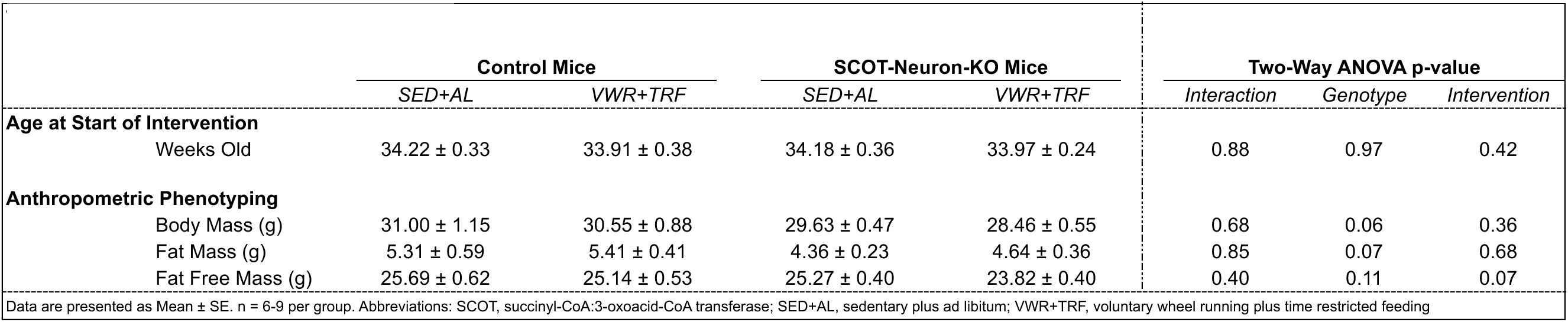
Male Baseline Anthropometric Data.

| Table S1. Male Baseline Anthropometric Data |  |  |  |  |  |  |  |
| --- | --- | --- | --- | --- | --- | --- | --- |
|  | Control Mice |  | SCOT-Neuron-KO Mice |  | Two-Way ANOVA p-value |  |  |
|  | SED+AL | VWR+TRF | SED+AL | VWR+TRF | Interaction | Genotype | Intervention |
| **Age at Start of Intervention** |  |  |  |  |  |  |  |
| Weeks Old | 34.22 ± 0.33 | 33.91 ± 0.38 | 34.18 ± 0.36 | 33.97 ± 0.24 | 0.88 | 0.97 | 0.42 |
| **Anthropometric Phenotyping** |  |  |  |  |  |  |  |
| Body Mass (g) | 31.00 ± 1.15 | 30.55 ± 0.88 | 29.63 ± 0.47 | 28.46 ± 0.55 | 0.68 | 0.06 | 0.36 |
| Fat Mass (g) | 5.31 ± 0.59 | 5.41 ± 0.41 | 4.36 ± 0.23 | 4.64 ± 0.36 | 0.85 | 0.07 | 0.68 |
| Fat Free Mass (g) | 25.69 ± 0.62 | 25.14 ± 0.53 | 25.27 ± 0.40 | 23.82 ± 0.40 | 0.40 | 0.11 | 0.07 |
| Data are presented as Mean ± SE. n = 6-9 per group. Abbreviations: SCOT, succinyl-CoA:3-oxoacid-CoA transferase; SED+AL, sedentary plus ad libitum; VWR+TRF, voluntary wheel running plus time restricted feeding |  |  |  |  |  |  |  |

## Acknowledgements and Funding

S10OD028598 (JPT), T32AG07811 (BAK), T32DK128770 (EF and SFS), T32HL144472 (KF), T32HL166142 (EDQ), K01DK112967 (EMM), P20GM144269 (EMM and JPT), R01AG069781 (JPT and PAC), and IDeA National Resource for Quantitative Proteomics and NIH/NIGMS grant R24GM137786.

## Author Contributions

Acquisition of funding (JPT, PAC), Investigation (SFS, BAK, EF, XD, FBB, JA, KLF, EDQ, EMM, PP), Supervision (JPT, PAC), Writing – Original Draft Preparation (SFS, JPT, PAC), Writing – Review & Editing (all authors)

## Statements and Declarations

The authors declare no competing interest.

## References

1. Hou Y, Dan X, Babbar M, Wei Y, Hasselbalch SG, Croteau DL, Bohr VA. Ageing as a risk factor for neurodegenerative disease. Nat Rev Neurol. 2019;15(10):565–81. Epub 20190909. doi: 10.1038/s41582-019-0244-7. PubMed PMID: 31501588.

2. Azam S, Haque ME, Balakrishnan R, Kim IS, Choi DK. The Ageing Brain: Molecular and Cellular Basis of Neurodegeneration. Front Cell Dev Biol. 2021;9:683459. Epub 20210813. doi: 10.3389/fcell.2021.683459. PubMed PMID: 34485280; PMCID: PMC8414981.

3. Lopez-Otin C, Blasco MA, Partridge L, Serrano M, Kroemer G. The hallmarks of aging. Cell. 2013;153(6):1194–217. doi: 10.1016/j.cell.2013.05.039. PubMed PMID: 23746838; PMCID: PMC3836174.

4. Guo J, Huang X, Dou L, Yan M, Shen T, Tang W, Li J. Aging and aging-related diseases: from molecular mechanisms to interventions and treatments. Signal Transduct Target Ther. 2022;7(1):391. Epub 20221216. doi: 10.1038/s41392-022-01251-0. PubMed PMID: 36522308; PMCID: PMC9755275.

5. Lopez-Otin C, Blasco MA, Partridge L, Serrano M, Kroemer G. Hallmarks of aging: An expanding universe. Cell. 2023;186(2):243–78. Epub 20230103. doi: 10.1016/j.cell.2022.11.001. PubMed PMID: 36599349.

6. Campisi J, Kapahi P, Lithgow GJ, Melov S, Newman JC, Verdin E. From discoveries in ageing research to therapeutics for healthy ageing. Nature. 2019;571(7764):183–92. Epub 20190710. doi: 10.1038/s41586-019-1365-2. PubMed PMID: 31292558; PMCID: PMC7205183.

7. Stessman J, Hammerman-Rozenberg R, Cohen A, Ein-Mor E, Jacobs JM. Physical activity, function, and longevity among the very old. Arch Intern Med. 2009;169(16):1476–83. doi: 10.1001/archinternmed.2009.248. PubMed PMID: 19752405.

8. Chakravarty EF, Hubert HB, Lingala VB, Fries JF. Reduced disability and mortality among aging runners: a 21-year longitudinal study. Arch Intern Med. 2008;168(15):1638–46. doi: 10.1001/archinte.168.15.1638. PubMed PMID: 18695077; PMCID: PMC3175643.

9. Wolinsky FD, Bentler SE, Hockenberry J, Jones MP, Obrizan M, Weigel PA, Kaskie B, Wallace RB. Long-term declines in ADLs, IADLs, and mobility among older Medicare beneficiaries. BMC Geriatr. 2011;11:43. Epub 20110816. doi: 10.1186/1471-2318-11-43. PubMed PMID: 21846400; PMCID: PMC3167753.

10. Colcombe SJ, Erickson KI, Raz N, Webb AG, Cohen NJ, McAuley E, Kramer AF. Aerobic fitness reduces brain tissue loss in aging humans. J Gerontol A Biol Sci Med Sci. 2003;58(2):176–80. doi: 10.1093/gerona/58.2.m176. PubMed PMID: 12586857.

11. Steiner JL, Murphy EA, McClellan JL, Carmichael MD, Davis JM. Exercise training increases mitochondrial biogenesis in the brain. J Appl Physiol (1985). 2011;111(4):1066–71. Epub 20110804. doi: 10.1152/japplphysiol.00343.2011. PubMed PMID: 21817111.

12. van Praag H, Kempermann G, Gage FH. Running increases cell proliferation and neurogenesis in the adult mouse dentate gyrus. Nat Neurosci. 1999;2(3):266–70. doi: 10.1038/6368. PubMed PMID: 10195220.

13. van Praag H, Christie BR, Sejnowski TJ, Gage FH. Running enhances neurogenesis, learning, and long-term potentiation in mice. Proc Natl Acad Sci U S A. 1999;96(23):13427–31. doi: 10.1073/pnas.96.23.13427. PubMed PMID: 10557337; PMCID: PMC23964.

14. He C, Sumpter R, Jr., Levine B. Exercise induces autophagy in peripheral tissues and in the brain. Autophagy. 2012;8(10):1548–51. Epub 20120815. doi: 10.4161/auto.21327. PubMed PMID: 22892563; PMCID: PMC3463459.

15. Mattson MP, Moehl K, Ghena N, Schmaedick M, Cheng A. Intermittent metabolic switching, neuroplasticity and brain health. Nat Rev Neurosci. 2018;19(2):63–80. Epub 20180111. doi: 10.1038/nrn.2017.156. PubMed PMID: 29321682; PMCID: PMC5913738.

16. Johnson RH, Walton JL, Krebs HA, Williamson DH. Post-exercise ketosis. Lancet. 1969;2(7635):1383-5. doi: 10.1016/s0140-6736(69)90931-3. PubMed PMID: 4188274.

17. Fery F, Balasse EO. Ketone body turnover during and after exercise in overnight-fasted and starved humans. Am J Physiol. 1983;245(4):E318–25. doi: 10.1152/ajpendo.1983.245.4.E318. PubMed PMID: 6353933.

18. Puchalska P, Crawford PA. Multi-dimensional Roles of Ketone Bodies in Fuel Metabolism, Signaling, and Therapeutics. Cell Metab. 2017;25(2):262–84. doi: 10.1016/j.cmet.2016.12.022. PubMed PMID: 28178565; PMCID: PMC5313038.

19. Puchalska P, Crawford PA. Metabolic and Signaling Roles of Ketone Bodies in Health and Disease. Annu Rev Nutr. 2021;41:49–77. doi: 10.1146/annurev-nutr-111120-111518. PubMed PMID: 34633859; PMCID: PMC8922216.

20. Fulghum K, Salathe SF, Davis X, Thyfault JP, Puchalska P, Crawford PA. Ketone body metabolism and cardiometabolic implications for cognitive health. NPJ Metab Health Dis. 2024;2. Epub 20241011. doi: 10.1038/s44324-024-00029-y. PubMed PMID: 40093558; PMCID: PMC11908690.

21. Venable AH, Lee LE, Feola K, Santoyo J, Broomfield T, Huen SC. Fasting-induced HMGCS2 expression in the kidney does not contribute to circulating ketones. Am J Physiol Renal Physiol. 2022;322(4):F460-F7. Epub 20220228. doi: 10.1152/ajprenal.00447.2021. PubMed PMID: 35224990; PMCID: PMC9076412.

22. Jensen NJ, Wodschow HZ, Nilsson M, Rungby J. Effects of Ketone Bodies on Brain Metabolism and Function in Neurodegenerative Diseases. Int J Mol Sci. 2020;21(22). Epub 20201120. doi: 10.3390/ijms21228767. PubMed PMID: 33233502; PMCID: PMC7699472.

23. Perez-Escuredo J, Van Hee VF, Sboarina M, Falces J, Payen VL, Pellerin L, Sonveaux P. Monocarboxylate transporters in the brain and in cancer. Biochim Biophys Acta. 2016;1863(10):2481–97. Epub 20160316. doi: 10.1016/j.bbamcr.2016.03.013. PubMed PMID: 26993058; PMCID: PMC4990061.

24. Cotter DG, Schugar RC, Wentz AE, d’Avignon DA, Crawford PA. Successful adaptation to ketosis by mice with tissue-specific deficiency of ketone body oxidation. Am J Physiol Endocrinol Metab. 2013;304(4):E363–74. Epub 20121211. doi: 10.1152/ajpendo.00547.2012. PubMed PMID: 23233542; PMCID: PMC3566508.

25. Fortier M, Castellano CA, Croteau E, Langlois F, Bocti C, St-Pierre V, Vandenberghe C, Bernier M, Roy M, Descoteaux M, Whittingstall K, Lepage M, Turcotte EE, Fulop T, Cunnane SC. A ketogenic drink improves brain energy and some measures of cognition in mild cognitive impairment. Alzheimers Dement. 2019;15(5):625–34. Epub 20190423. doi: 10.1016/j.jalz.2018.12.017. PubMed PMID: 31027873.

26. Shippy DC, Wilhelm C, Viharkumar PA, Raife TJ, Ulland TK. beta-Hydroxybutyrate inhibits inflammasome activation to attenuate Alzheimer’s disease pathology. J Neuroinflammation. 2020;17(1):280. Epub 20200921. doi: 10.1186/s12974-020-01948-5. PubMed PMID: 32958021; PMCID: PMC7507727.

27. Cunnane S, Nugent S, Roy M, Courchesne-Loyer A, Croteau E, Tremblay S, Castellano A, Pifferi F, Bocti C, Paquet N, Begdouri H, Bentourkia M, Turcotte E, Allard M, Barberger-Gateau P, Fulop T, Rapoport SI. Brain fuel metabolism, aging, and Alzheimer’s disease. Nutrition. 2011;27(1):3–20. Epub 20101029. doi: 10.1016/j.nut.2010.07.021. PubMed PMID: 21035308; PMCID: PMC3478067.

28. Marosi K, Kim SW, Moehl K, Scheibye-Knudsen M, Cheng A, Cutler R, Camandola S, Mattson MP. 3-Hydroxybutyrate regulates energy metabolism and induces BDNF expression in cerebral cortical neurons. J Neurochem. 2016;139(5):769–81. Epub 20161114. doi: 10.1111/jnc.13868. PubMed PMID: 27739595; PMCID: PMC5123937.

29. Haces ML, Hernandez-Fonseca K, Medina-Campos ON, Montiel T, Pedraza-Chaverri J, Massieu L. Antioxidant capacity contributes to protection of ketone bodies against oxidative damage induced during hypoglycemic conditions. Exp Neurol. 2008;211(1):85–96. Epub 20080126. doi: 10.1016/j.expneurol.2007.12.029. PubMed PMID: 18339375.

30. Finn PF, Dice JF. Ketone bodies stimulate chaperone-mediated autophagy. J Biol Chem. 2005;280(27):25864–70. Epub 20050509. doi: 10.1074/jbc.M502456200. PubMed PMID: 15883160.

31. Evans M, Cogan KE, Egan B. Metabolism of ketone bodies during exercise and training: physiological basis for exogenous supplementation. J Physiol. 2017;595(9):2857–71. Epub 20161207. doi: 10.1113/JP273185. PubMed PMID: 27861911; PMCID: PMC5407977.

32. Queathem ED, Moazzami Z, Stagg DB, Nelson AB, Fulghum K, Hayir A, Seay A, Gillingham JR, d’Avignon DA, Han X, Ruan HB, Crawford PA, Puchalska P. Ketogenesis supports hepatic polyunsaturated fatty acid homeostasis via fatty acid elongation. Sci Adv. 2025;11(5):eads0535. Epub 20250129. doi: 10.1126/sciadv.ads0535. PubMed PMID: 39879309; PMCID: PMC11777252.

33. Chadman KK, Gong S, Scattoni ML, Boltuck SE, Gandhy SU, Heintz N, Crawley JN. Minimal aberrant behavioral phenotypes of neuroligin-3 R451C knockin mice. Autism Res. 2008;1(3):147–58. doi: 10.1002/aur.22. PubMed PMID: 19360662; PMCID: PMC2701211.

34. Martin-Montalvo A, Mercken EM, Mitchell SJ, Palacios HH, Mote PL, Scheibye-Knudsen M, Gomes AP, Ward TM, Minor RK, Blouin MJ, Schwab M, Pollak M, Zhang Y, Yu Y, Becker KG, Bohr VA, Ingram DK, Sinclair DA, Wolf NS, Spindler SR, Bernier M, de Cabo R. Metformin improves healthspan and lifespan in mice. Nat Commun. 2013;4:2192. doi: 10.1038/ncomms3192. PubMed PMID: 23900241; PMCID: PMC3736576.

35. Stover KR, Campbell MA, Van Winssen CM, Brown RE. Early detection of cognitive deficits in the 3xTg-AD mouse model of Alzheimer’s disease. Behav Brain Res. 2015;289:29–38. Epub 20150417. doi: 10.1016/j.bbr.2015.04.012. PubMed PMID: 25896362.

36. Berta Sunyer SP, Harald Höger, Gert Lubec. Barnes maze, a useful task to assess spatial reference memory in the mice. Protocol Exchange. 2007. doi: 10.1038/nprot.2007.390.

37. Morris EM, Noland RD, Allen JA, McCoin CS, Xia Q, Koestler DC, Shook RP, Lighton JRB, Christianson JA, Thyfault JP. Difference in Housing Temperature-Induced Energy Expenditure Elicits Sex-Specific Diet-Induced Metabolic Adaptations in Mice. Obesity (Silver Spring). 2020;28(10):1922–31. Epub 20200828. doi: 10.1002/oby.22925. PubMed PMID: 32857478; PMCID: PMC7511436.

38. Franczak E, Maurer A, Drummond VC, Kugler BA, Wells E, Wenger M, Peelor FF, 3rd, Crosswhite A, McCoin CS, Koch LG, Britton SL, Miller BF, Thyfault JP. Divergence in aerobic capacity and energy expenditure influence metabolic tissue mitochondrial protein synthesis rates in aged rats. Geroscience. 2024;46(2):2207–22. Epub 20231026. doi: 10.1007/s11357-023-00985-1. PubMed PMID: 37880490; PMCID: PMC10828174.

39. Searle BC, Pino LK, Egertson JD, Ting YS, Lawrence RT, MacLean BX, Villen J, MacCoss MJ. Chromatogram libraries improve peptide detection and quantification by data independent acquisition mass spectrometry. Nat Commun. 2018;9(1):5128. Epub 20181203. doi: 10.1038/s41467-018-07454-w. PubMed PMID: 30510204; PMCID: PMC6277451.

40. Graw S, Tang J, Zafar MK, Byrd AK, Bolden C, Peterson EC, Byrum SD. proteiNorm - A User-Friendly Tool for Normalization and Analysis of TMT and Label-Free Protein Quantification. ACS Omega. 2020;5(40):25625–33. Epub 20200930. doi: 10.1021/acsomega.0c02564. PubMed PMID: 33073088; PMCID: PMC7557219.

41. Thurman TJ WC, Alkam D, Bird JT, Gies A, Dhusia K, Robeson II MS, and Byrum SD. proteoDA: a package for quantitive proteomics. Journal of Open Source Software. 2023. doi: 10.21105/joss.05184.

42. Bolstad B. preprocessCore: A collection of pre=processing functions2019.

43. Ritchie ME, Phipson B, Wu D, Hu Y, Law CW, Shi W, Smyth GK. limma powers differential expression analyses for RNA-sequencing and microarray studies. Nucleic Acids Res. 2015;43(7):e47. Epub 20150120. doi: 10.1093/nar/gkv007. PubMed PMID: 25605792; PMCID: PMC4402510.

44. Chawade A, Alexandersson E, Levander F. Normalyzer: a tool for rapid evaluation of normalization methods for omics data sets. J Proteome Res. 2014;13(6):3114–20. Epub 20140502. doi: 10.1021/pr401264n. PubMed PMID: 24766612; PMCID: PMC4053077.

45. Huber W, von Heydebreck A, Sultmann H, Poustka A, Vingron M. Variance stabilization applied to microarray data calibration and to the quantification of differential expression. Bioinformatics. 2002;18 Suppl 1:S96–104. doi: 10.1093/bioinformatics/18.suppl_1.s96. PubMed PMID: 12169536.

46. Alhamdoosh M, Ng M, Wilson NJ, Sheridan JM, Huynh H, Wilson MJ, Ritchie ME. Combining multiple tools outperforms individual methods in gene set enrichment analyses. Bioinformatics. 2017;33(3):414–24. doi: 10.1093/bioinformatics/btw623. PubMed PMID: 27694195; PMCID: PMC5408797.

47. Bolstad BM, Irizarry RA, Astrand M, Speed TP. A comparison of normalization methods for high density oligonucleotide array data based on variance and bias. Bioinformatics. 2003;19(2):185–93. doi: 10.1093/bioinformatics/19.2.185. PubMed PMID: 12538238.

48. Puopolo T, Seeram NP, Liu C. Chloroform/Methanol Protein Extraction and In-solution Trypsin Digestion Protocol for Bottom-up Proteomics Analysis. Bio Protoc. 2024;14(16):e5055. Epub 20240820. doi: 10.21769/BioProtoc.5055. PubMed PMID: 39210950; PMCID: PMC11349489.

49. Franczak E, Kugler BA, Salathe SF, Allen JA, Sardiu ME, McCoin CS, Hevener AL, Morris EM, Thyfault JP. Loss of ovarian function prevents exercise-induced activation of hepatic mitophagic flux. Am J Physiol Endocrinol Metab. 2025;328(6):E869-E84. Epub 20250428. doi: 10.1152/ajpendo.00107.2025. PubMed PMID: 40293097; PMCID: PMC12148014.

50. d’Avignon DA, Puchalska P, Ercal B, Chang Y, Martin SE, Graham MJ, Patti GJ, Han X, Crawford PA. Hepatic ketogenic insufficiency reprograms hepatic glycogen metabolism and the lipidome. JCI Insight. 2018;3(12). Epub 20180621. doi: 10.1172/jci.insight.99762. PubMed PMID: 29925686; PMCID: PMC6124396.

51. Hughey CC, Bracy DP, Rome FI, Goelzer M, Donahue EP, Viollet B, Foretz M, Wasserman DH. Exercise training adaptations in liver glycogen and glycerolipids require hepatic AMP-activated protein kinase in mice. Am J Physiol Endocrinol Metab. 2024;326(1):E14-E28. Epub 20231108. doi: 10.1152/ajpendo.00289.2023. PubMed PMID: 37938177; PMCID: PMC11193517.

52. Sharma A, Oonthonpan L, Sheldon RD, Rauckhorst AJ, Zhu Z, Tompkins SC, Cho K, Grzesik WJ, Gray LR, Scerbo DA, Pewa AD, Cushing EM, Dyle MC, Cox JE, Adams C, Davies BS, Shields RK, Norris AW, Patti G, Zingman LV, Taylor EB. Impaired skeletal muscle mitochondrial pyruvate uptake rewires glucose metabolism to drive whole-body leanness. Elife. 2019;8. Epub 20190718. doi: 10.7554/eLife.45873. PubMed PMID: 31305240; PMCID: PMC6684275.

53. Edmond J, Robbins RA, Bergstrom JD, Cole RA, de Vellis J. Capacity for substrate utilization in oxidative metabolism by neurons, astrocytes, and oligodendrocytes from developing brain in primary culture. J Neurosci Res. 1987;18(4):551–61. doi: 10.1002/jnr.490180407. PubMed PMID: 3481403.

54. Takashima N, Odaka YS, Sakoori K, Akagi T, Hashikawa T, Morimura N, Yamada K, Aruga J. Impaired cognitive function and altered hippocampal synapse morphology in mice lacking Lrrtm1, a gene associated with schizophrenia. PLoS One. 2011;6(7):e22716. Epub 20110727. doi: 10.1371/journal.pone.0022716. PubMed PMID: 21818371; PMCID: PMC3144940.

55. Soler-Llavina GJ, Arstikaitis P, Morishita W, Ahmad M, Sudhof TC, Malenka RC. Leucine-rich repeat transmembrane proteins are essential for maintenance of long-term potentiation. Neuron. 2013;79(3):439–46. doi: 10.1016/j.neuron.2013.06.007. PubMed PMID: 23931994; PMCID: PMC3741667.

56. Dufort-Gervais J, Provost C, Charbonneau L, Norris CM, Calon F, Mongrain V, Brouillette J. Neuroligin-1 is altered in the hippocampus of Alzheimer’s disease patients and mouse models, and modulates the toxicity of amyloid-beta oligomers. Sci Rep. 2020;10(1):6956. Epub 20200424. doi: 10.1038/s41598-020-63255-6. PubMed PMID: 32332783; PMCID: PMC7181681.

57. Kim J, Park D, Seo NY, Yoon TH, Kim GH, Lee SH, Seo J, Um JW, Lee KJ, Ko J. LRRTM3 regulates activity-dependent synchronization of synapse properties in topographically connected hippocampal neural circuits. Proc Natl Acad Sci U S A. 2022;119(3). doi: 10.1073/pnas.2110196119. PubMed PMID: 35022233; PMCID: PMC8784129.

58. Bhouri M, Morishita W, Temkin P, Goswami D, Kawabe H, Brose N, Sudhof TC, Craig AM, Siddiqui TJ, Malenka R. Deletion of LRRTM1 and LRRTM2 in adult mice impairs basal AMPA receptor transmission and LTP in hippocampal CA1 pyramidal neurons. Proc Natl Acad Sci U S A. 2018;115(23):E5382-E9. Epub 20180521. doi: 10.1073/pnas.1803280115. PubMed PMID: 29784826; PMCID: PMC6003336.

59. de Wit J, Sylwestrak E, O’Sullivan ML, Otto S, Tiglio K, Savas JN, Yates JR, 3rd, Comoletti D, Taylor P, Ghosh A. LRRTM2 interacts with Neurexin1 and regulates excitatory synapse formation. Neuron. 2009;64(6):799–806. doi: 10.1016/j.neuron.2009.12.019. PubMed PMID: 20064388; PMCID: PMC2829666.

60. Sudhof TC. Neuroligins and neurexins link synaptic function to cognitive disease. Nature. 2008;455(7215):903–11. doi: 10.1038/nature07456. PubMed PMID: 18923512; PMCID: PMC2673233.

61. Bertin F, Moya-Alvarado G, Quiroz-Manriquez E, Ibacache A, Kohler-Solis A, Oliva C, Sierralta J. Dlg Is Required for Short-Term Memory and Interacts with NMDAR in the Drosophila Brain. Int J Mol Sci. 2022;23(16). Epub 20220816. doi: 10.3390/ijms23169187. PubMed PMID: 36012453; PMCID: PMC9409279.

62. Guzman M, Blazquez C. Is there an astrocyte-neuron ketone body shuttle? Trends Endocrinol Metab. 2001;12(4):169–73. doi: 10.1016/s1043-2760(00)00370-2. PubMed PMID: 11295573.

63. Varela L, Kim JG, Fernandez-Tussy P, Aryal B, Liu ZW, Fernandez-Hernando C, Horvath TL. Astrocytic lipid metabolism determines susceptibility to diet-induced obesity. Sci Adv. 2021;7(50):eabj2814. Epub 20211210. doi: 10.1126/sciadv.abj2814. PubMed PMID: 34890239; PMCID: PMC11323787.

64. MoTr PACSG, Lead A, MoTr PACSG. Temporal dynamics of the multi-omic response to endurance exercise training. Nature. 2024;629(8010):174–83. Epub 20240501. doi: 10.1038/s41586-023-06877-w. PubMed PMID: 38693412; PMCID: PMC11062907.

65. Schonfeld P, Reiser G. Why does brain metabolism not favor burning of fatty acids to provide energy? Reflections on disadvantages of the use of free fatty acids as fuel for brain. J Cereb Blood Flow Metab. 2013;33(10):1493–9. Epub 20130807. doi: 10.1038/jcbfm.2013.128. PubMed PMID: 23921897; PMCID: PMC3790936.

66. Thevenet J, De Marchi U, Domingo JS, Christinat N, Bultot L, Lefebvre G, Sakamoto K, Descombes P, Masoodi M, Wiederkehr A. Medium-chain fatty acids inhibit mitochondrial metabolism in astrocytes promoting astrocyte-neuron lactate and ketone body shuttle systems. FASEB J. 2016;30(5):1913–26. Epub 20160202. doi: 10.1096/fj.201500182. PubMed PMID: 26839375.

67. Auestad N, Korsak RA, Morrow JW, Edmond J. Fatty acid oxidation and ketogenesis by astrocytes in primary culture. J Neurochem. 1991;56(4):1376–86. doi: 10.1111/j.1471-4159.1991.tb11435.x. PubMed PMID: 2002348.

68. Blazquez C, Sanchez C, Velasco G, Guzman M. Role of carnitine palmitoyltransferase I in the control of ketogenesis in primary cultures of rat astrocytes. J Neurochem. 1998;71(4):1597–606. doi: 10.1046/j.1471-4159.1998.71041597.x. PubMed PMID: 9751193.

69. Sonnay S, Chakrabarti A, Thevenet J, Wiederkehr A, Christinat N, Masoodi M. Differential Metabolism of Medium-Chain Fatty Acids in Differentiated Human-Induced Pluripotent Stem Cell-Derived Astrocytes. Front Physiol. 2019;10:657. Epub 20190604. doi: 10.3389/fphys.2019.00657. PubMed PMID: 31214043; PMCID: PMC6558201.

70. Le Foll C, Dunn-Meynell AA, Miziorko HM, Levin BE. Regulation of hypothalamic neuronal sensing and food intake by ketone bodies and fatty acids. Diabetes. 2014;63(4):1259–69. Epub 20131230. doi: 10.2337/db13-1090. PubMed PMID: 24379353; PMCID: PMC3964505.

71. Le Foll C, Dunn-Meynell AA, Miziorko HM, Levin BE. Role of VMH ketone bodies in adjusting caloric intake to increased dietary fat content in DIO and DR rats. Am J Physiol Regul Integr Comp Physiol. 2015;308(10):R872–8. Epub 20150318. doi: 10.1152/ajpregu.00015.2015. PubMed PMID: 25786485; PMCID: PMC4436979.

72. Le Foll C, Levin BE. Fatty acid-induced astrocyte ketone production and the control of food intake. Am J Physiol Regul Integr Comp Physiol. 2016;310(11):R1186–92. Epub 20160427. doi: 10.1152/ajpregu.00113.2016. PubMed PMID: 27122369; PMCID: PMC4935491.

