## Supplementary figures and images for "Neuronal ketone body utilization couples exercise and time-restricted feeding to cognitive enhancement"

### Supplemental Figure 1

#### Male SCOT-Neuron-KO

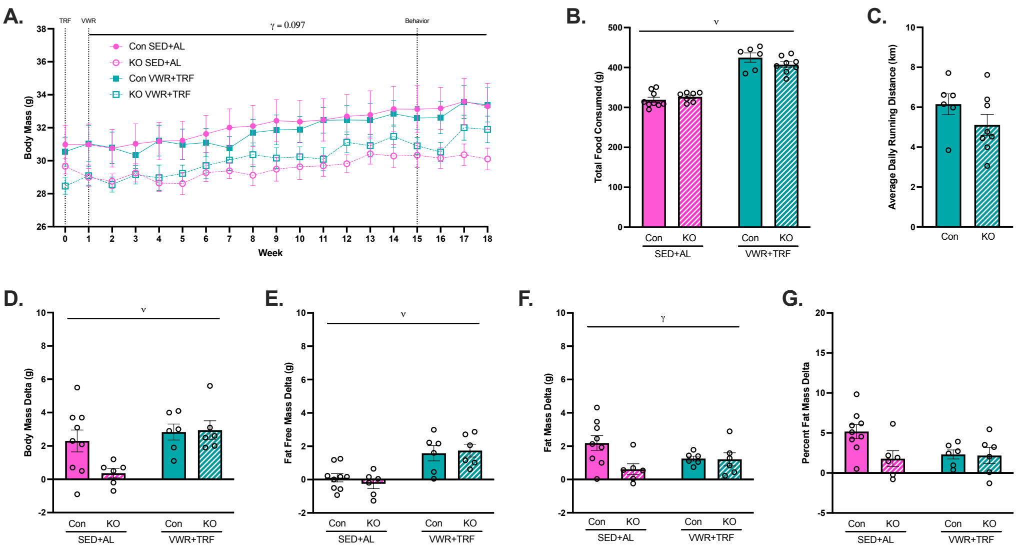

### Supplemental Figure 2

#### Male SCOT-Neuron-KO

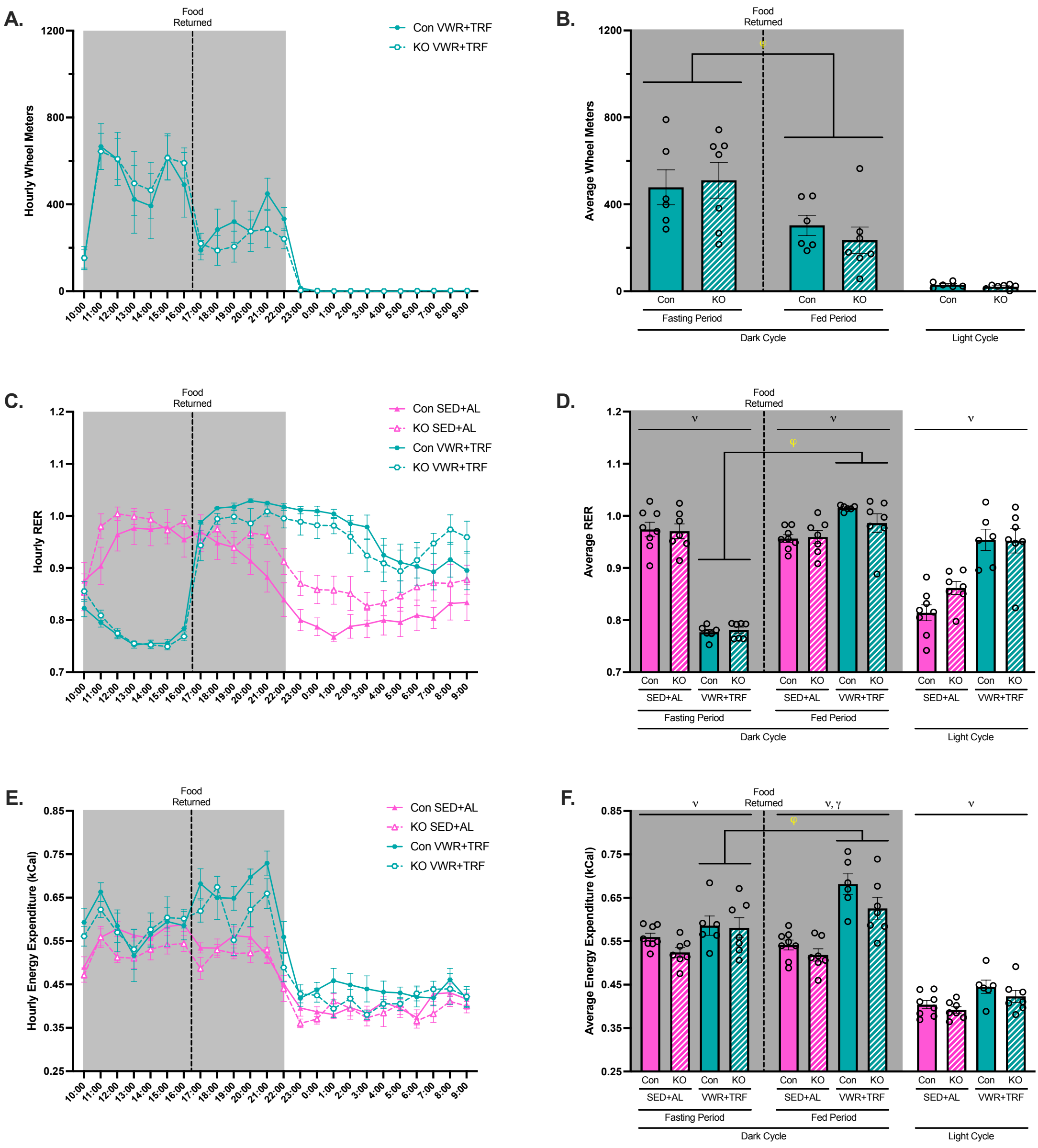

### Supplemental Figure 3

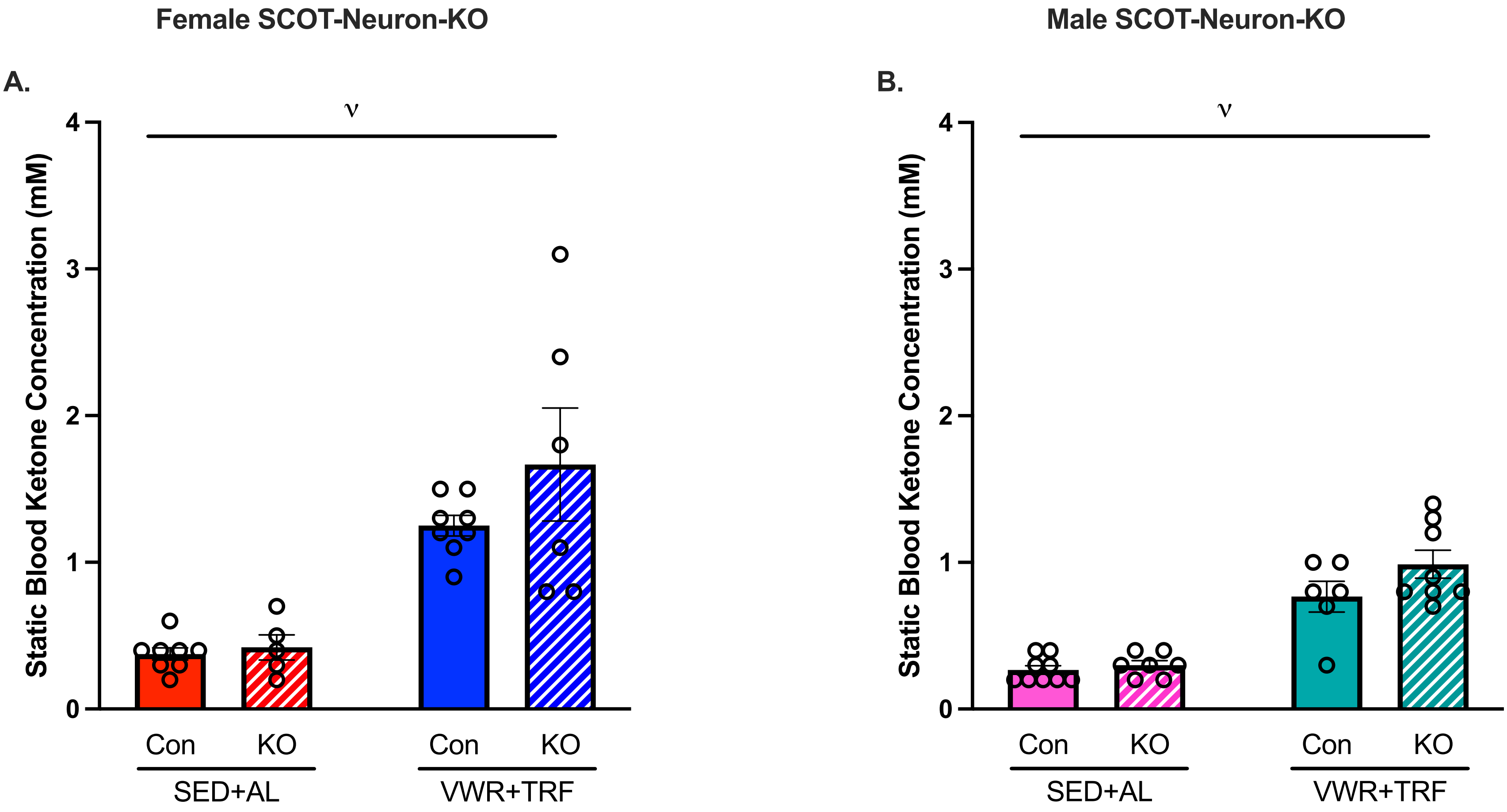

### Supplemental Figure 4

#### Male SCOT-Neuron-KO

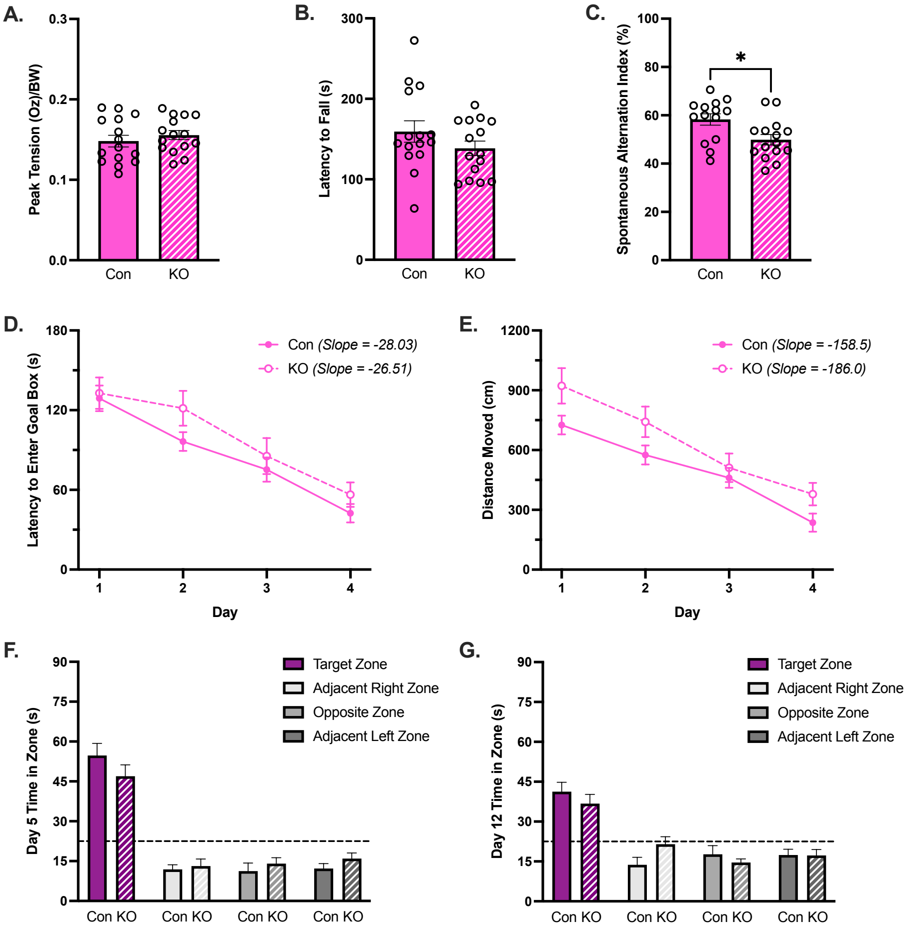

### Supplemental Figure 5

#### Male SCOT-Neuron-KO

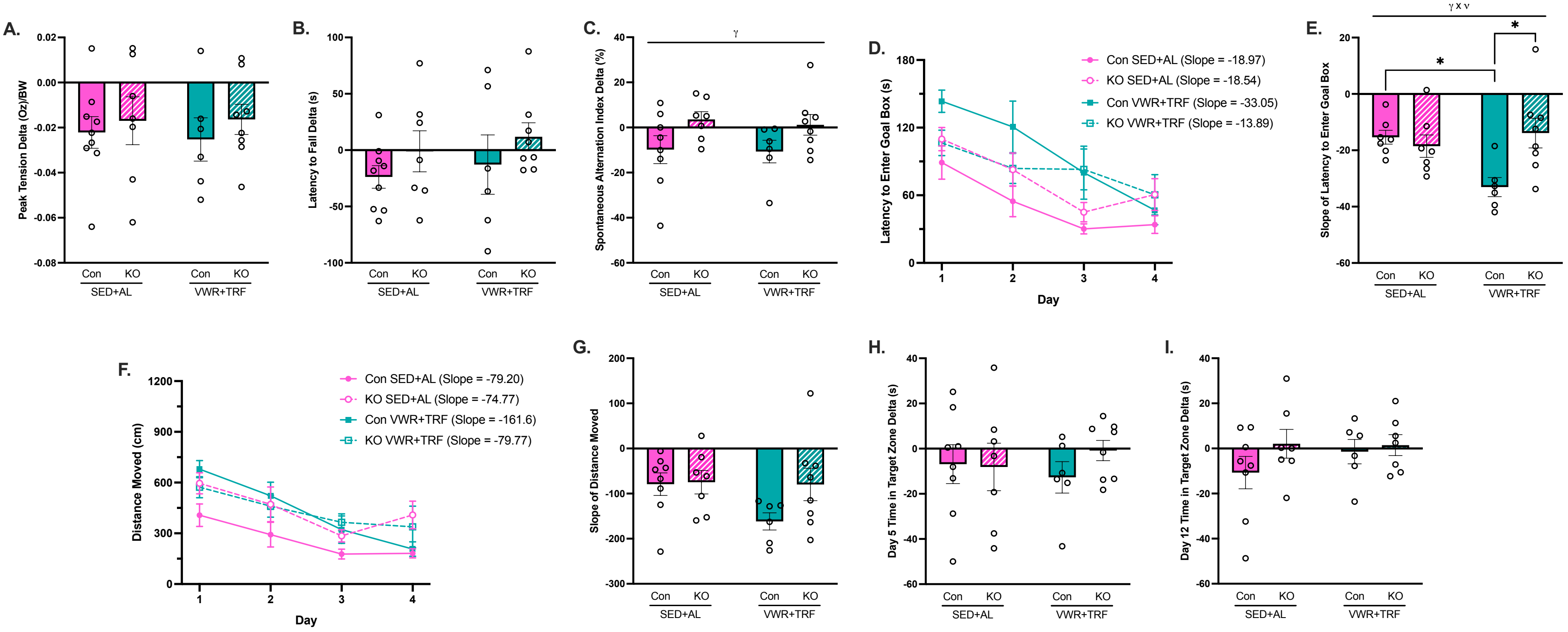

### Supplemental Figure 6

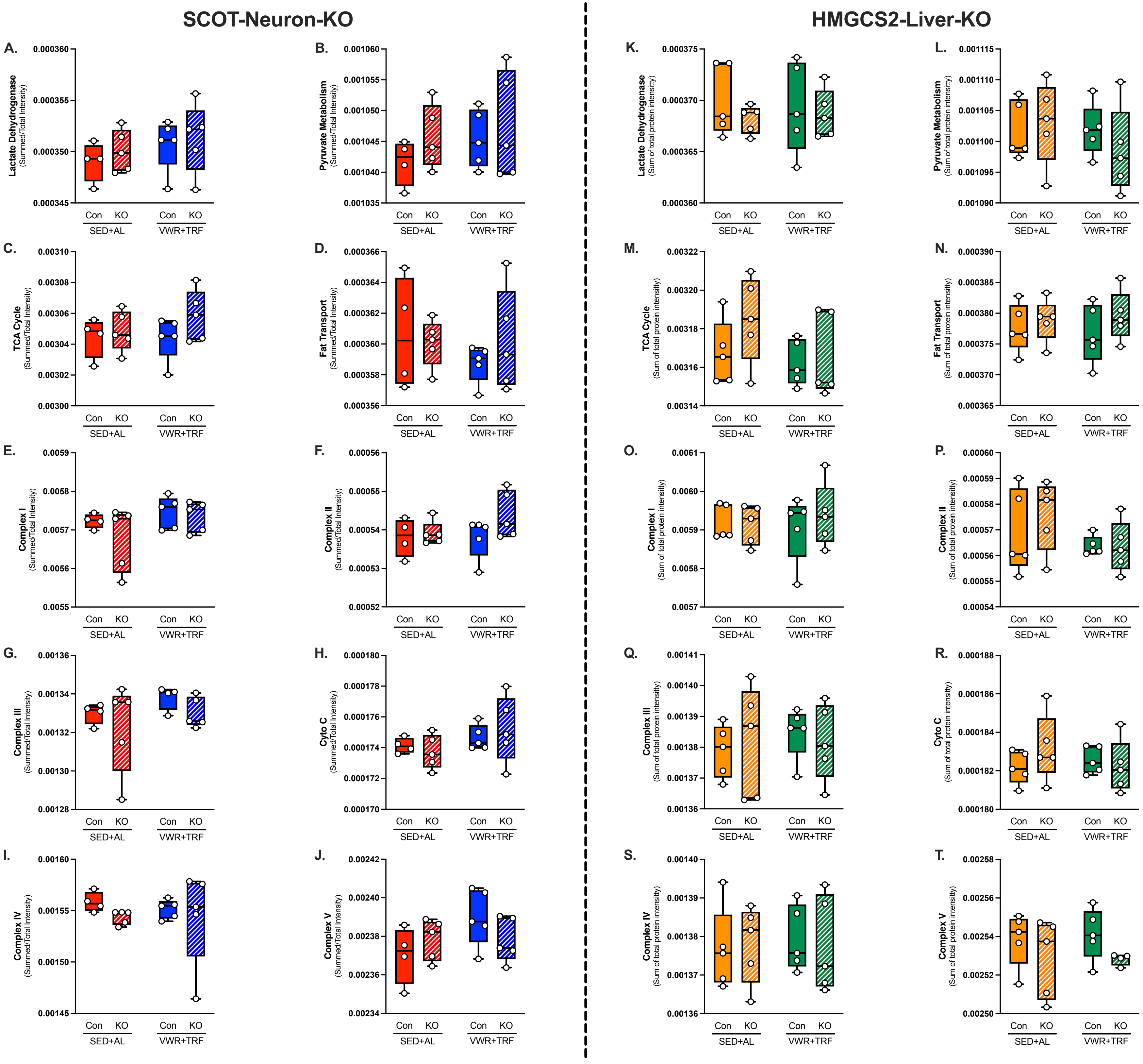

### Supplemental Figure 7

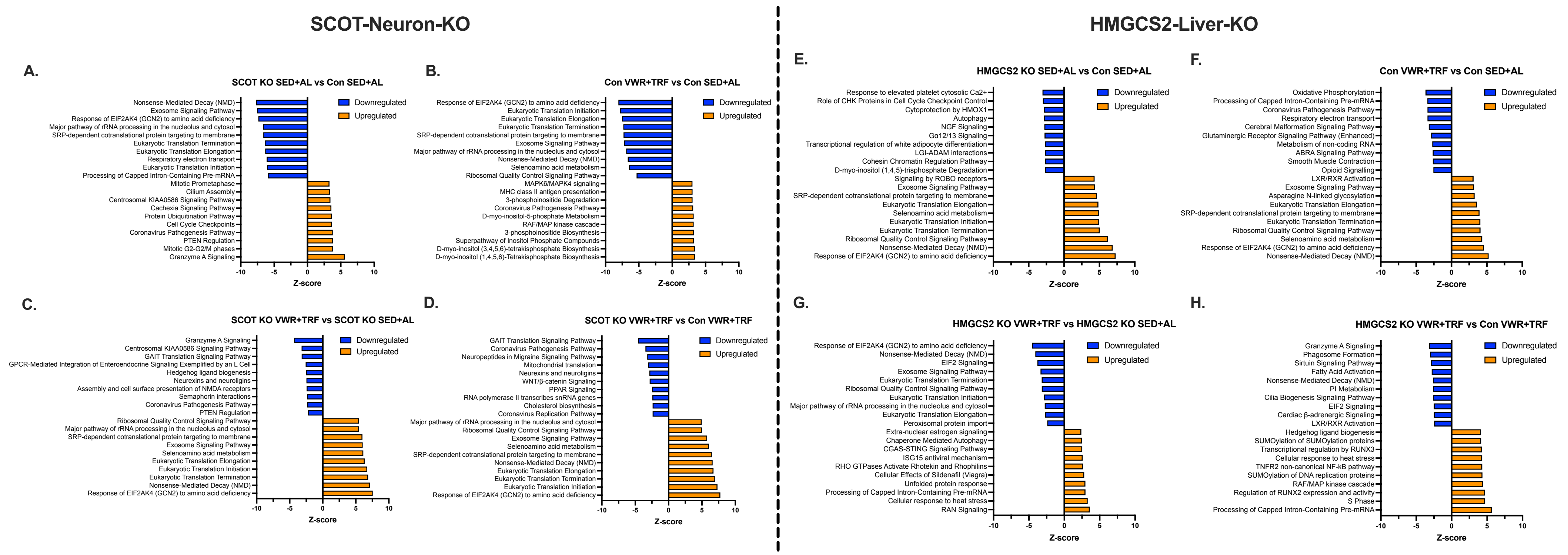
